# ProLigGPT: a language model for protein-targeted ligand design free from structural constraints

**DOI:** 10.1101/2023.06.29.543848

**Authors:** Yuesen Li, Chengyi Gao, Xin Song, Xiangyu Wang, Xinyue Tan, Wenqiang Ma, Andy Peng Xiang, Kai Xia, Hong Chen, Hang Fan, Fei Wang, Jianan Li, Yutiantian Lei, Hemeng Zhang, Yihan Wang, Jinlu Ma, Yiping Li, Yungang Xu, Suxia Han

## Abstract

The deep-learning language model offers promising approaches to molecular design. However, existing models typically exhibit one or more of the following limitations: structural constraints, confinement to a single task, and lack of cross-domain capability (e.g., protein-to-ligand generation). Inspired by GPT’s success in language modeling, we introduce ProLigGPT, a seq2seq autoregressive model based on GPT-2 and trained on ∼1.9 million protein-ligand pairs without 3D structures. ProLigGPT supports tasks including *de novo* ligand generation and scaffold-based molecular design for target proteins. Benchmarking against three state-of-the-art structure-based models across eight representative target proteins, ProLigGPT demonstrated superior performance on comprehensive metrics. Through systematic evaluations of ProLigGPT’s interpretability, we find that ProLigGPT uses the full protein sequence to generate target-aware ligands, unlike pocket-based strategies that depend only on residues near the binding site. Applying ProLigGPT to Pl3Kα inhibitor design successfully yielded inhibitors with a novel scaffold and inhibitors derived from a known scaffold, demonstrating that ProLigGPT can reach beyond the boundaries of known chemical space. Finally, we validated the generalization ability of ProLigGPT on Target T1, which lacks a crystal structure and known inhibitors, confirming that ProLigGPT can design inhibitors solely from sequence information. These findings highlight the promise of language models in designing ligands for target proteins free from structural constraints, offering a flexible and scalable framework for exploring vast chemical spaces in drug discovery.

## Introduction

Drug discovery has long been a cornerstone of therapeutic innovation, driving advances in the treatment of a wide range of diseases. Over the past decades, computational approaches have profoundly accelerated this process^1^. Early computer-aided drug design (CADD) methods, including molecular docking^2^, quantitative structure–activity relationships^3^ (QSAR), and pharmacophore modeling^4^, provided efficient strategies for virtual screening and candidate prioritization. However, the immense size of the drug-like chemical space, estimated to be up to 10^60^ molecules^5^, renders exhaustive exploration infeasible with CADD approaches, underscoring the need for more powerful computational paradigms.

The emergence of artificial intelligence for drug design (AIDD) has reshaped the field by integrating target protein and compound information for virtual screening and molecular design. AIDD methods leverage diverse molecular and protein representations, spanning one-dimensional (1D) sequences, two-dimensional (2D) graphs and contact maps, and three-dimensional (3D) conformations. In these frameworks, compounds may be represented as SMILES^6^ (Simplified Molecular Input Line Entry System) strings, molecular fingerprints^7^, molecular graphs, or chemical structures, while proteins can be described by amino acid sequences, residue–residue contact maps^8^, or folded structures. Deep learning architectures, including convolutional neural networks^9^, recurrent neural networks^10^, variational autoencoders^11^, generative adversarial networks^12^, diffusion models^13^, and contrastive learning^14^ (e.g., CLIP), have been applied to these representations, enabling significant progress in virtual screening and molecular generation^15^. In particular, target-aware ligand generation, which uses target protein information to directly generate potentially active molecules and moves beyond exhaustive searches of known compounds, has become an important direction in AIDD.

Most existing target-aware ligand generation methods follow a structure-based drug design (SBDD) approach: they assume the availability of experimentally resolved or high-confidence predicted protein 3D structures, typically together with predefined binding pockets. In real ligand design scenarios, this assumption is often not satisfied. First, a large fraction of proteins still lack reliable experimental structures or high-confidence predicted structures. Previous statistics showed that, among 14,849 known protein families, fewer than one-third had experimentally solved structures^16^. In the human reference proteome, only 1,663 proteins (9.93%) have more than 90% of their amino acid sequences covered by experimentally resolved PDB structures; even when high-confidence AlphaFold structures are included (pLDDT ≥ 70 and coverage ≥ 85%), only 4,930 human proteins (26.79%) meet this criterion^17^. This means that more than two-thirds of human protein targets cannot be directly addressed by conventional structure-dependent ligand design. Second, even when a protein structure is available, accurate small-molecule binding-site prediction remains an unresolved challenge. AF2BIND^18^ explicitly notes that identifying ligandable sites remains an outstanding problem, especially for de novo small-molecule binding-site prediction. Third, current structure-based drug discovery workflows often depend on high-quality holo structures or conformationally stable apo pocket structures with clear boundaries, making them highly sensitive to structure quality, pocket conformational stability, and preprocessing accuracy^19^. Therefore, structure-based molecular generation is largely restricted to targets with high-quality structures and well-defined binding pockets. This limitation is particularly pronounced for newly identified or understudied targets, including many disease-associated proteins, membrane proteins, and functionally uncharacterized proteins. For many of these targets, reliable structural models and binding-site annotations are unavailable, with few known small-molecule ligands. By contrast, protein sequences are widely available and, like natural languages, are information-complete^20^. Their amino acid order efficiently stores structural and functional information. This disparity raises a fundamental question: can chemically valid, drug-like candidate ligands be generated for a given target from its amino acid sequence alone, without using its 3D structure or a predefined binding pocket?

The evolution of Transformer-based models offers a potential route to addressing this question. With attention mechanisms at its core, the Transformer has been widely adopted as a foundational architecture for modern language models^21^; continued advances in attention design have broadened the range of natural language processing (NLP) tasks that these models can address while steadily improving their performance^22,23^. Moreover, proteins and molecules, both naturally expressible as sequences, are well aligned with the paradigm of large-scale language modeling. Building on these advances, we hypothesize that language models can learn bioactivity relationships from paired protein-ligand sequences, enabling the direct generation of candidate ligands from a target protein sequence without 3D structures or predefined binding pockets. This sequence-based formulation not only simplifies the design pipeline from “sequence → structure → pocket → molecule” to “sequence → molecule” but also enables systematic evaluation of whether language models can extract sufficient biochemical information solely from sequence data to guide precise and efficient molecular design.

In practice, Transformer-based models differ markedly in their suitability for molecular design, reflecting variations in attention design and architecture. Transformer architectures are classified into three types: 1) encoder-decoder models (e.g., standard Transformer^21^), mainly used for translation; 2) encoder-only models (e.g., BERT^22^), designed for contextual understanding; and 3) decoder-only models (e.g., GPT^23^), primarily for text generation. The representative models for molecular design with transformer architectures are summarized in Table S1 (Supplementary Materials Part 1). Encoder-decoder models, such as Grechishnikova’s model^24^, Uludoğan’s model^25^, and AlphaDrug^26^, enable the decoder to interact with encoder outputs via cross-attention, while treating ligand generation as a direct translation from protein sequences to molecular sequences. Although effective for mapping protein “language” to chemical “language,” these models are limited in flexibility and struggle to incorporate complex conditional constraints, making it difficult to perform multitask generation, such as scaffold-based ligand design. Encoder-only models, which exclusively rely on bidirectional self-attention and lack a decoder module, are fundamentally unsuitable for molecular design tasks as they cannot perform generative tasks or conditionally control the generation process. In contrast, decoder-only models, inspired by the GPT family^23,27^, employ causal self-attention to approach molecule design as an autoregressive generation task. By conditioning on tags or property descriptors, models such as MolGPT^28^ and ProGen^29^ enable multitask generation and property control. However, these models are restricted to single-domain conditioning (e.g., molecular properties or protein annotations) and thus lack explicit target-awareness, that is, the ability to design ligands specifically tailored to protein targets. In addition, several Transformer-based molecular design models rely on structural information, such as AlphaDrug, TamGen^30^, and PrefixMol^31^, which limits their generalizability and makes them unsuitable for settings where only the protein sequence is available.

To this end, we present ProLigGPT, a sequence-based autoregressive generative model for small-molecule design. Built upon a decoder-only Transformer architecture, ProLigGPT is trained in two stages: first, it learns the chemical language from SMILES strings; second, it incorporates protein-ligand pairs to capture activity relationships between proteins and compounds. Using Byte Pair Encoding^32^ (BPE) tokenization, the model achieves efficient joint representation of proteins and molecules, facilitating cross-domain learning. Using only sequences as conditional inputs, ProLigGPT achieves three innovations simultaneously: 1) independence from structural data (“free from structure”), ensuring applicability across the proteome regardless of structural availability; 2) target-aware ligand generation, enabling design specifically guided by the protein sequence; and 3) multitask capability, supporting both *de novo* design and scaffold-based design. By leveraging the autoregressive generation mechanism, ProLigGPT can either directly explore novel chemical space or decorate known scaffolds through their corresponding ligand prompts, which is a flexible strategy particularly well-suited for scaffold-based ligand design.

We evaluated the performance of ProLigGPT through both *in silico* and wet-lab validations on well-studied targets as well as less well-characterized targets. In computational experiments, comparative evaluations were conducted on eight protein targets (including four low-homology targets not present in the ProLigGPT training set) against structure-based models such as PocketFlow^33^, Lingo3DMol^34^, and TargetDiff^13^. ProLigGPT exhibited superior performance in key metrics such as quantitative estimate of drug-likeness^35^ (QED), synthetic accessibility^36^ (SA), and docking scores^37^. Notably, ProLigGPT achieved state-of-the-art (SOTA) results in a comprehensive assessment of these key metrics. This advantage was further corroborated on the TarPass^38^ benchmark, which comprises 20 protein structures representative of real-world drug discovery settings and 15 baseline models spanning non-3D, 3D in situ generative, and optimization-based approaches. On the well-characterized protein PI3Kα, wet-lab experiments confirmed ProLigGPT’s ability to generate compounds with novel scaffolds and to design derivatives based on known inhibitor scaffolds. Both classes of molecules were successfully synthesized and exhibited significant biological activity. This experimental validation confirmed the practicality of ProLigGPT-designed molecules in terms of chemical validity, synthesizability, and biological activity. It further showed that ProLigGPT can access chemical space beyond that covered by existing molecules and scaffolds, thereby expanding target-relevant bioactive chemical space. For Target T1, which lacks any reported crystal structure or inhibitors, ProLigGPT achieved a structure-free approach to ligand design using only the target protein sequence. The model’s zero-shot generalization to this unseen protein target was experimentally validated through wet-lab assays, which identified four of the generated candidates as inhibitors. To our knowledge, this is the first demonstration that small-molecule inhibitors can be designed directly from the protein sequence of a structurally unresolved GPCR and experimentally validated, with neither molecule generation nor candidate selection involving structural modeling, binding-pocket definition, or molecular docking. Overall, ProLigGPT provides a simple and practical sequence-driven molecular design platform that generates candidate ligand libraries using the target protein’s amino acid sequence as its sole target-specific input. This direct “sequence → ligand” design paradigm does not rely on the target structure, predefined binding-pocket information, or reference ligands, thereby extending small-molecule design to understudied protein targets.

## Results

### An overview of the framework of ProLigGPT

ProLigGPT is a sequence-to-sequence (seq2seq) autoregressive model built on the generative pre-trained transformer, designed to generate molecules that target a specific protein. Its architecture is tailored to prioritize molecular bioactivity during molecule generation, enabling it to efficiently propose candidate compounds from protein sequence data. The model was trained on a large-scale dataset comprising approximately 1.9 million protein-ligand pairs (Methods, Supplementary Materials Part 2), which provides experimentally validated binding affinities but lacks protein-ligand complex structures^39^. By relying on this extensive experimental dataset, ProLigGPT addresses the limitations of structure-based models, allowing it to generate molecules that reflect bioactivity with the target protein rather than their structural interactions. Unlike structure-based models, ProLigGPT employs a modular framework, decoupling the direct modeling of protein-ligand interactions from molecule generation. Detailed protein-ligand interactions, while critical, are evaluated post hoc using molecular docking and related computational methods. This decouple ensures that the generating process remains focused on achieving favorable bioactivity, while downstream analyses refine the molecular interactions.

ProLigGPT supports two molecular design modes: 1) *de novo* design, using only the protein sequence to generate entirely novel molecular structures, thereby exploring untapped chemical space (Fig.1a, Opt1); 2) scaffold-based design, incorporating both the protein sequence and the SMILES representation of a known ligand scaffold (ligand prompt strategy), generates derivative molecules that are guided by the chemical essence of the input scaffold while extending known chemical spaces (Fig.1a, Opt2). The selected design mode dictates the input text for the generating process. In the generating process, the ProLigGPT tokenizer (Methods, Supplementary Materials Part 3) first converts the input text into tokens and encodes them as input IDs for the model (Fig.1b–d). The ProLigGPT model (Methods, Supplementary Materials Part 4) then autoregressively predicts the sequence of token IDs representing the generated molecule. Once the sequence is generated, the tokenizer decodes the IDs into SMILES strings, the standard representations of chemical structures (Fig.1e–f). To ensure the generated molecules are chemically valid and stable, post-processing (Methods) is conducted using Open Babel^40^ to convert the SMILES strings into SDF format and perform energy minimization, enabling further analysis and practical applications (Fig.1g–h). In summary, ProLigGPT offers a reliable and flexible framework for generating bioactive molecules, with a workflow that emphasizes efficiency, chemical diversity, and bioactivity-driven design.

**Fig.1.**
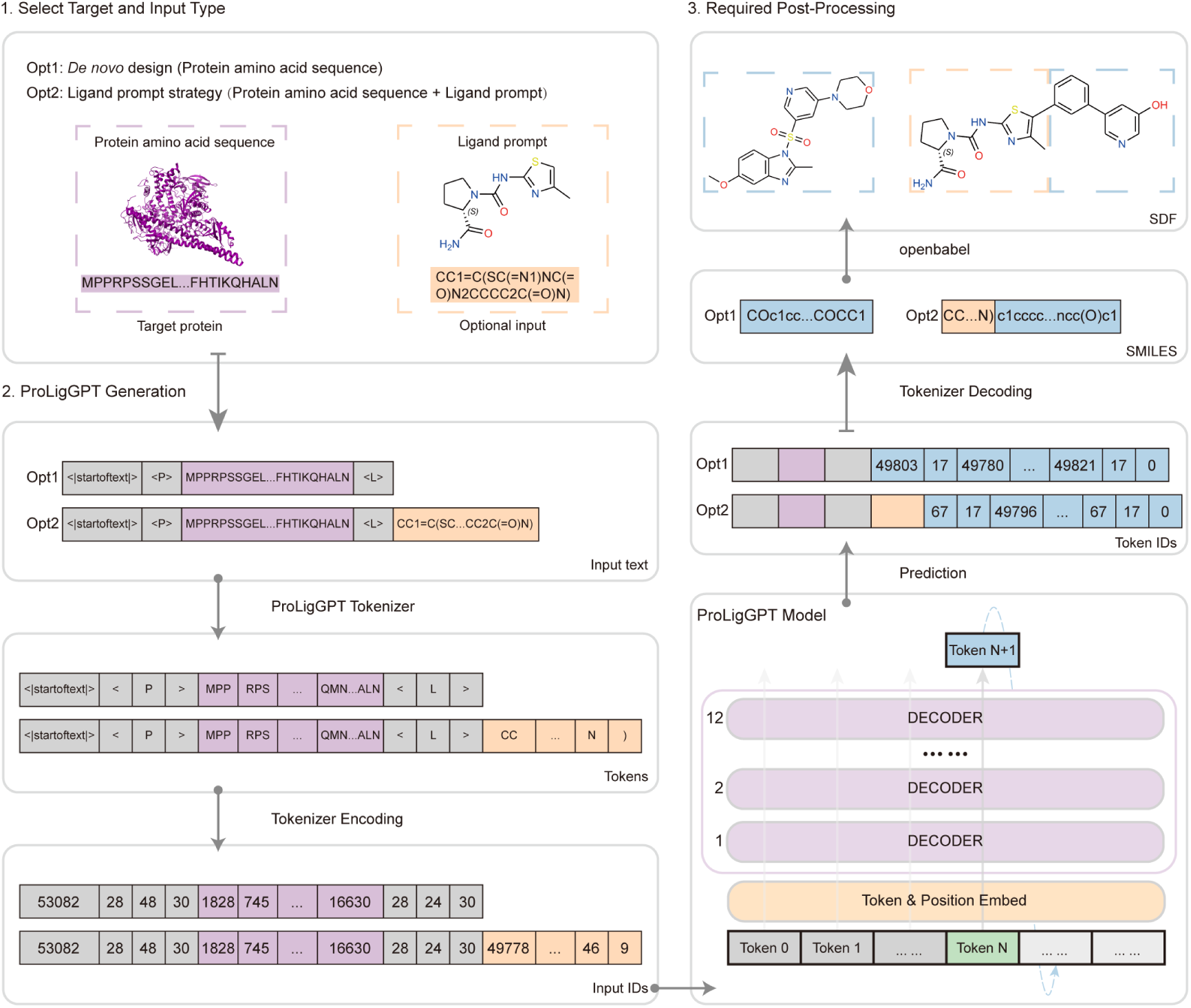
The schematic view and generative process of ProLigGPT. **a,** ProLigGPT takes inputs with two options, either only protein sequence (Opt1, for *de novo* ligand design) or protein sequence with the SMILES of a known ligand scaffold (Opt2, for generation based on the ligand prompt strategy). **b–d,** ProLigGPT tokenizer encodes the input sequences of protein and ligand SMILES into token IDs for the model training and generation. **e,** ProLigGPT model autoregressively predicts the ligand represented as a sequence of token IDs. **f–g,** ProLigGPT tokenizer decodes the generated tokens into the SMILES of the generated molecules. **h,** The post-processing step involves using Open Babel to convert the SMILES into SDF format, followed by energy minimization to ensure the generated structures are both chemically stable and viable.

### ProLigGPT excels in designing drug-like molecules with favorable docking results

To evaluate the performance of ProLigGPT, we compared it with three SOTA structure-based drug design (SBDD) models, including PocketFlow, Lingo3DMol, and TargetDiff. Eight diverse proteins were selected for this comparative evaluation: PI3Kα (PDB: 4jps), HAT1 (PDB: 6vo5), YTHDC1 (PDB: 4r3i), FAK1 (PDB: 3bz3), BRAF (PDB: 3d4q), GRK4 (PDB: 4yhj), LONP1 (PDB: 7oxo), and SARM1 (PDB: 7cm6). PI3Kα was the primary focus of this study, while HAT1, YTHDC1, FAK1, BRAF, and GRK4 were previously reported in the literature associated with the compared SBDD models. Additionally, HAT1, YTHDC1, LONP1, and SARM1 had no significant homologous proteins in the ProLigGPT training dataset, as confirmed by BLAST analysis, with maximum sequence identities below 25%. For each target protein, we generated 10,000 molecules per model using the default settings (Supplementary Materials Part 5). The generated molecules were evaluated based on molecular properties and metrics, including Vina score^41^, QED, SA, logP^42^, TPSA^43^, molecular weight, atom count, and SMILES length (Supplementary Materials Part 6), as summarized in Table 1. The distribution of these properties and metrics illustrates the performance variability among different models and target proteins (Extended Data Fig.1). When comparing models on PI3Kα, ProLigGPT showed comparable performance across the SA, QED, and Vina score metrics. However, evaluating model performance based on a single metric may be influenced by each model’s inherent tendencies during molecule generation. For instance, as shown in Extended Data Fig.1, PocketFlow tends to generate smaller molecules, leading to lower SA values; in contrast, Lingo3DMol tends to favor larger molecules, which may yield better Vina score but perform worse on QED (Supplementary Materials Part 7). Across evaluations of all target proteins, ProLigGPT consistently performed well, achieving the best Vina score in general, and ranking highly on QED.

**Table 1.** The evaluation of models for de novo design across different target proteins. For each model, we generated 10,000 molecules using the default settings for each of the eight target proteins. ^a^Four proteins, HAT1, YTHDC1, LONP1, and SARM1, were analyzed using BLAST and had maximum sequence identities of 21.9%, 19.7%, 19.7%, and 20.8%, respectively, to proteins in the ProLigGPT training set. Vina score, QED, and SA are shown as mean values. The bold values indicate the best performance results across various target proteins. **Valid generation,** the number of molecules generated by the model that could be successfully converted into pdbqt format for molecular docking. **Vina score,** a scoring function from AutoDock Vina, evaluates binding affinity (kcal/mol), where more negative values indicate stronger potential interactions between the ligand and the target protein. **QED,** Quantitative Estimate of Drug-likeness, a drug-likeness index that varies from 0 (least drug-like) to 1 (most drug-like). **SA,** Synthetic Accessibility, an index reflecting the synthetic difficulty of molecules, ranging from 0 (easy to synthesize) to 10 (extremely difficult to synthesize). **Unique,** The percentage of unique molecules among the total generated molecules. **Drug-like targeted molecules,** defined using a comprehensive metric for drug-like molecules with a QED score of at least 0.3, an SA score of no more than 5, and filtered to remove duplicate molecules, additionally require a Vina score of -6.5 or lower to ensure favorable binding affinity with the target protein.

|  | **Model** | **Valid generation** | **Vina score ↓** | **QED ↑** | **SA ↓** | **Unique, %↑** | **Drug-like target molecules ↑** |
|---|---|---|---|---|---|---|---|
| PI3Kα | ProLigGPT | 9997 | -9.014 | 0.575 | 2.965 | 99.19% | **9776** |
|  | PocketFlow | 9996 | -7.881 | 0.506 | **2.339** | 69.81% | 5055 |
|  | Lingo3DMol | 10356 | **-9.274** | 0.197 | 3.868 | 97.24% | 1185 |
|  | TargetDiff | 9574 | -7.613 | **0.649** | 4.774 | **99.96%** | 4210 |
| HAT1<sup>a</sup> | ProLigGPT | 9630 | **-9.189** | **0.609** | 2.943 | 99.44% | **8933** |
|  | PocketFlow | 9990 | -8.692 | 0.465 | **2.575** | 82.13% | 5143 |
|  | Lingo3DMol | 10014 | -9.122 | 0.382 | 4.255 | 77.88% | 4554 |
|  | TargetDiff | 9076 | -8.827 | 0.527 | 4.736 | **99.97%** | 4808 |
| YTHDC1<sup>a</sup> | ProLigGPT | 9917 | **-7.347** | 0.560 | **2.472** | **98.54%** | **7518** |
|  | PocketFlow | 9975 | -6.400 | **0.585** | 2.886 | 72.84% | 3550 |
|  | Lingo3DMol | 9999 | -6.169 | 0.450 | 3.823 | 44.05% | 2258 |
|  | TargetDiff | 9469 | -5.780 | 0.479 | 4.227 | 98.27% | 901 |
| FAK1 | ProLigGPT | 9985 | **-9.265** | 0.531 | 2.586 | 99.22% | **9739** |
|  | PocketFlow | 9989 | -7.776 | 0.540 | **2.447** | 67.66% | 4996 |
|  | Lingo3DMol | 10030 | -8.102 | 0.359 | 4.126 | 83.74% | 3799 |
|  | TargetDiff | 9260 | -7.866 | **0.623** | 4.684 | **99.92%** | 4733 |
| BRAF | ProLigGPT | 9962 | **-10.345** | 0.475 | **2.624** | 99.21% | **9323** |
|  | PocketFlow | 9990 | -9.067 | **0.553** | 2.667 | 81.24% | 6176 |
|  | Lingo3DMol | 10018 | -10.232 | 0.254 | 3.634 | 78.05% | 2198 |
|  | TargetDiff | 9519 | -9.347 | 0.493 | 5.207 | **99.90%** | 3229 |
| GRK4 | ProLigGPT | 9904 | **-8.915** | **0.558** | 2.777 | 99.49% | **9667** |
|  | PocketFlow | 9987 | -8.036 | 0.498 | **2.464** | 72.41% | 4861 |
|  | Lingo3DMol | 10014 | -8.664 | 0.290 | 4.002 | 93.00% | 3255 |
|  | TargetDiff | 9495 | -8.669 | 0.553 | 5.210 | **99.94%** | 3722 |
| LONP1<sup>a</sup> | ProLigGPT | 9743 | **-8.151** | 0.549 | **2.608** | 99.35% | **8719** |
|  | PocketFlow | 9992 | -7.007 | **0.577** | 2.687 | 77.26% | 5193 |
|  | Lingo3DMol | 10133 | -8.053 | 0.309 | 4.338 | 84.85% | 3134 |
|  | TargetDiff | 8010 | -7.307 | 0.334 | 5.546 | **99.96%** | 760 |
| SARM1<sup>a</sup> | ProLigGPT | 9799 | **-7.138** | **0.541** | **2.784** | 98.95% | **6791** |
|  | PocketFlow | 9981 | -5.417 | 0.446 | 3.433 | 80.57% | 1021 |
|  | Lingo3DMol | 10017 | -6.936 | 0.227 | 4.485 | 75.24% | 744 |
|  | TargetDiff | 7522 | -6.654 | 0.221 | 5.723 | **99.93%** | 417 |

Given these observations, it is crucial to comprehensively evaluate drug candidates by integrating multiple metrics, as relying solely on a single metric (such as the Vina score) may result in a biased assessment. While some molecules achieve favorable Vina scores, their potential as drug candidates could be limited by low QED values or poor synthetic feasibility^34^. Therefore, we used a comprehensive evaluation strategy that includes QED, SA, and Vina score to identify promising drug candidates. As illustrated in Extended Data Fig.2, molecules generated by each model were filtered through four sequential criteria: Vina score ≤ -6.5, QED ≥ 0.3, and SA ≤ 5, followed by deduplication, to select drug-like targeted molecules. This filtering process is aimed at ensuring that the generated molecules are not only predicted to bind well to the target proteins but are also synthetically feasible and exhibit desirable drug-like properties. Following this filtering process, ProLigGPT showed SOTA performance in generating drug-like targeted molecules, demonstrating its substantial potential for advancing drug design applications. Further benchmarking with TarPass^38^, which compares models from non-3D, 3D in situ and optimization-based generation paradigms across representative targets from diverse protein classes, showed that ProLigGPT still retained the largest total number of candidates after multi-stage screening (Supplementary Materials Part 8).

These findings highlight a critical challenge in drug design, wherein relying solely on structural information does not guarantee the generation of molecules simultaneously possessing high biological activity, desirable synthetic accessibility (SA), and ideal medicinal chemistry properties (QED). The experimentally validated structures typically utilized by SBDD models are limited, and docking-based structures often fail to accurately reflect genuine interactions, thus compromising model effectiveness. In contrast, our study suggests that reducing dependency on structural information and effectively leveraging abundant high-quality experimental data facilitates the generation of candidate molecules with desirable drug-like characteristics. Consequently, sequence-based ligand design methods offer notable advantages, including greater flexibility and the effective use of extensive datasets, providing robust and efficient solutions for drug discovery.

### ProLigGPT exploits the full protein sequence for molecular design and exhibits sensitivity to point mutations

ProLigGPT is a language model that solely relies on protein sequence information, eschewing the processing of structural data or pocket information. This approach provides a clear research paradigm to explore how language models utilize protein sequences for effective molecular design. To facilitate this, we developed an interactive Colab notebook that explicitly visualizes the model’s attention distributions (Methods). Users can intuitively observe ProLigGPT’s decision-making processes by inputting a target protein sequence along with the corresponding molecule generated by ProLigGPT. Using PI3Kα as an illustrative example, ProLigGPT comprehensively utilizes each token within the protein sequence to achieve molecular design. This approach significantly differs from pocket-based methods, which specifically concentrate on particular amino acid residues that directly interact with ligands.

Upon converting protein amino acid sequences into token sequences, the information carried by each token depends on both its internal amino acid composition and the sequential order of the tokens within the sequence. To validate this, we analyzed the token compositions and their characteristics, as well as compared molecules generated from original sequences against those from sequences with shuffled input token order (Supplementary Materials Part 9). Statistical analysis of token frequency in the training set revealed that tokens containing four amino acids were the most common (Fig.S8a). Furthermore, using AlphaFold3^44^, we predicted structures for tokens longer than ten amino acids from the 3000 most frequent tokens, finding high confidence in these structural predictions (Fig.S8b,c). We then conducted comparative analyses using four protein sequences (HAT1, YTHDC1, LONP1, and SARM1), selected for their lack of significant homology with sequences in ProLigGPT’s training dataset. Each protein sequence was tested both in its original form and after shuffling the order of its input tokens. The outcomes of these comparisons, summarized in Table 2, demonstrated a significant deterioration in predicted binding affinity (higher Vina scores) for molecules generated from sequences with shuffled tokens. These molecules also exhibited reduced desirability, as reflected by lower QED scores and worsened SA values. Distribution patterns of these metrics also shifted noticeably, showing increased standard deviations and a preference for generating smaller molecules (Fig.S9–12). These observations indicate that shuffling token sequences impairs ProLigGPT’s accurate interpretation of protein sequence information, increasing confusion and causing the model to adopt a more conservative generation strategy favoring smaller molecules. These findings reinforce that ProLigGPT effectively utilizes comprehensive protein sequence information, with token sequence arrangement playing a pivotal role in molecular design.

**Table 2.** Comparison of ProLigGPT-generated molecules using original and shuffled tokens. a. using original tokens as input. b. using shuffled tokens as input. Data are presented as mean ± SD. Bolded mean values indicate superior performance for metrics, while bolded SD values highlight greater variability. For the three key molecular metrics (Vina score, QED, and SA), normal tokens generally yielded better results, except for a marginally higher QED observed for shuffled tokens in SARM1. Molecules generated using shuffled tokens exhibited altered physicochemical properties, including decreased logP and increased TPSA. Regarding molecular weight, atom count (excluding hydrogens), and SMILES length, ProLigGPT generated molecules with lower atom counts when using shuffled tokens as input. The SD values for all molecular properties and metrics were higher when shuffled tokens were used.

|  |  | Vina score ↓ | QED ↑ | SA ↓ | LogP | TPSA | Molecular weight | Atom count | SMILES length |
|---|---|---|---|---|---|---|---|---|---|
| HAT1 | a | **-9.189** ± 1.119 | **0.609** ± 0.187 | **2.943** ± 0.685 | 2.561 ± 2.051 | 95.914 ± 41.267 | 386.166 ± 76.890 | 27.115 ± 5.309 | 53.725 ± 12.215 |
|  | b | -8.262 ± **1.182** | 0.457 ± **0.206** | 3.777 ± **0.803** | -0.400 ± **2.427** | 136.360 ± **55.775** | 354.823 ± **98.248** | 24.558 ± **6.555** | 60.508 ± **17.424** |
| YTHDC1 | a | **-7.347** ± 0.968 | **0.560** ± 0.164 | **2.472** ± 0.538 | 3.588 ± 1.510 | 83.583 ± 34.727 | 362.570 ± 73.359 | 26.029 ± 5.083 | 46.238 ± 10.685 |
|  | b | -6.841 ± **1.032** | 0.557 ± **0.206** | 3.103 ± **1.012** | 1.242 ± **2.647** | 108.693 ± **54.692** | 337.193 ± **87.455** | 23.712 ± **5.939** | 51.083 ± **17.121** |
| LONP1 | a | **-8.151** ± 0.892 | **0.549** ± 0.176 | **2.608** ± 0.652 | 3.400 ± 2.039 | 88.938 ± 38.255 | 381.774 ± 76.639 | 26.838 ± 5.228 | 46.238 ± 10.685 |
|  | b | -7.447 ± **1.138** | 0.488 ± **0.213** | 3.432 ± **0.999** | 0.281 ± **2.826** | 122.454 ± **57.863** | 340.435 ± **96.258** | 23.706 ± **6.559** | 51.083 ± **17.121** |
| SARM1 | a | **-7.138** ± 0.861 | 0.541 ± 0.194 | **2.784** ± 0.784 | 2.764 ± 2.174 | 97.525 ± 48.769 | 358.807 ± 81.628 | 25.325 ± 5.426 | 48.643 ± 14.087 |
|  | b | -6.756 ± **0.897** | **0.559** ± **0.206** | 3.139 ± **0.968** | 1.049 ± **2.578** | 109.693 ± **52.531** | 333.932 ± **90.225** | 23.351 ± **6.121** | 51.081 ± **16.766** |

As a sequence-based model, ProLigGPT needs to sensitively detect and respond to significant point mutations. We validated this by examining the PI3Kα H1047R mutation^45^, where histidine at position 1047 is replaced by arginine (Supplementary Materials Part 10). We generated 10,000 *de novo* molecules separately using the wild-type PI3Kα sequence (UniProt ID: P42336) and the H1047R mutant sequence. The resulting molecular distributions differed significantly between these two inputs (Fig.S13), and some molecules generated in response to the mutation exhibited a notable tendency toward allosteric pockets^46^ at the docking level (Fig.S14). Additionally, by preserving the scaffold of STX-478^47^, we preliminarily indicated the potential of ProLigGPT’s scaffold-based design mode (ligand prompt strategy) in responding to mutated sequences (Fig.S15).

In conclusion, ProLigGPT effectively leverages comprehensive protein sequence information for molecular design and possesses sensitivity to sequence variations. This capability fundamentally distinguishes ProLigGPT from traditional pocket-based approaches, as it exploits global protein sequence information rather than focusing solely on local residues directly involved in ligand interactions.

### ProLigGPT innovates ligands with novel scaffolds

For proteins with abundant known ligands, if the model only captures and learns frequent patterns from these ligand molecules, the generated molecules will become overly conservative, making it difficult to break away from existing molecular patterns in the training set and limiting exploration into broader chemical spaces. To empirically assess the implications of the issue outlined above in practical drug design, we selected PI3Kα, a key member of the PI3K family extensively studied in drug discovery, as a representative case. PI3Kα plays a crucial role in regulating cellular growth, proliferation, and survival, thus serving as a significant therapeutic target for various solid tumors^48,49^. We performed wet-lab experiments to evaluate ProLigGPT’s ability to *de novo* design small-molecule ligands with novel scaffolds that inhibit PI3Kα activity (Supplementary Materials Parts 5, 11, and 12).

The *de novo* design mode of ProLigGPT (Fig.1a, Opt1) was used to generate 100,000 potential ligands, relying solely on the protein sequence of PI3Kα (Fig.2a, Methods). Specifically, the generated SMILES and their 3D structures were saved individually in 100,000 SDF files. After energy minimization, 7 structurally invalid SDF files were discarded. The 99,993 remaining ligands fall into three categories as shown in Fig.2b: 1) ‘Known ligand recall’ includes 2,552 ligands from the training dataset with known inhibitory activity against PI3Kα (Extended Data Fig.3a); 2) ‘Ligand repurposing’ refers to 1,201 ligands originally developed to target other proteins but identified by ProLigGPT as potential inhibitors of PI3Kα (Extended Data Fig.3b); 3) ‘Innovative molecules’ includes 96,240 molecules not present in existing ligand databases and without documented interactions with proteins (Extended Data Fig.3c), showcasing ProLigGPT’s ability to explore novel chemical spaces beyond current drug repositories. The generated ligands exhibit molecular properties highly similar to those of known PI3Kα ligands, demonstrating the model’s ability to effectively capture the essential characteristics of these known ligands (Extended Data Fig.4).

**Fig.2.**
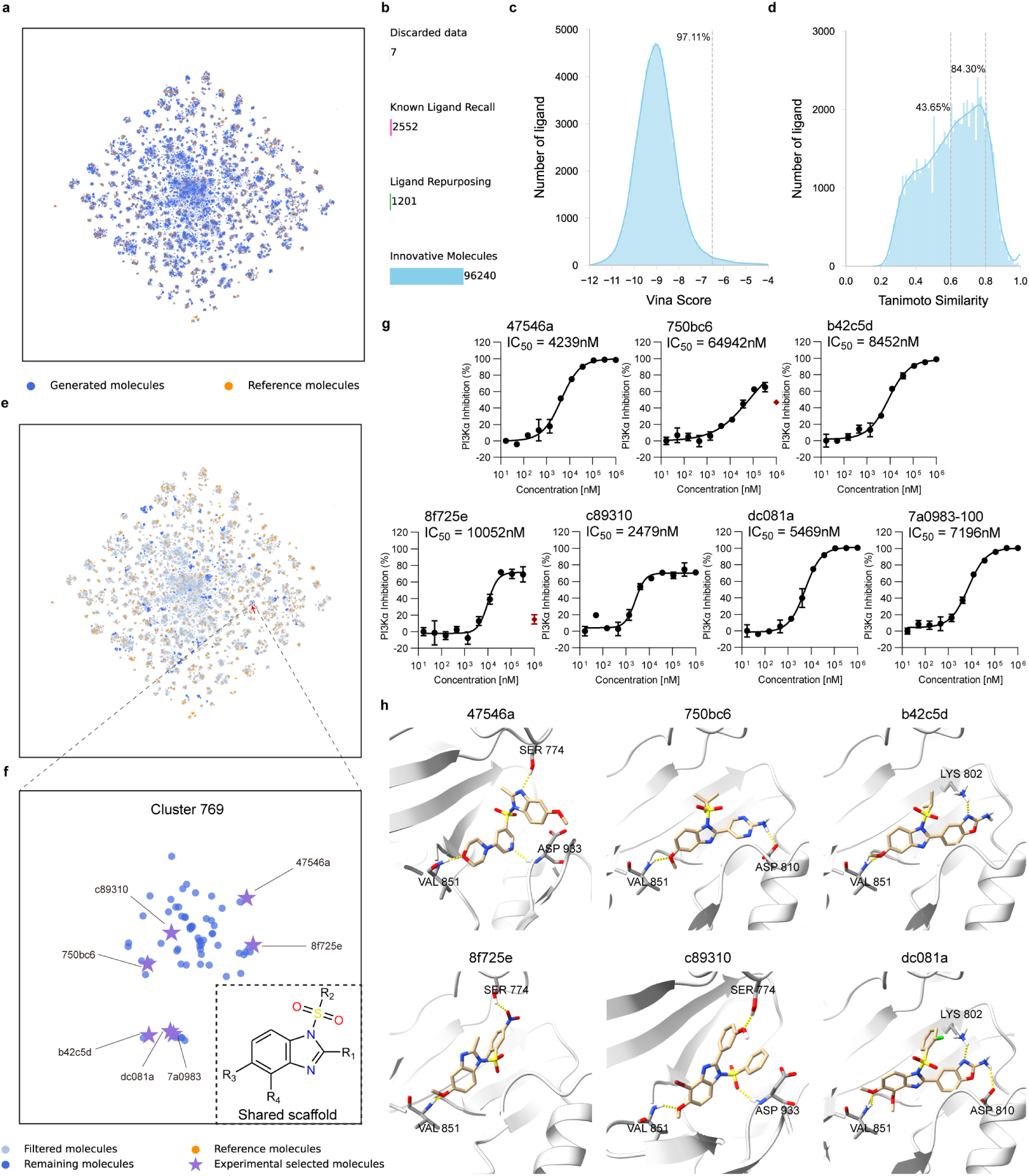
Discovery of PI3K*α* ligands through *de novo* design. **a,** t-SNE visualization of generated molecules (royalblue points) and reference molecules (darkorange points). **b,** Classification of generated molecules. **c,** Frequency distribution histogram of Vina score for innovative molecules. **d,** Frequency distribution histogram of the Tanimoto similarity for innovative molecules. The value represents the maximum Tanimoto similarity between the innovative molecule and all PI3Kα ligands in the training set. **e–f,** Using the Louvain method, clusters containing reference molecules (lightsteelblue points) were filtered out. Among the remaining clusters, Cluster 769 received special attention, with its molecules sharing the 1-(sulfonyl)-1*H*-benzimidazole scaffold. Seven molecules from this cluster were selected for experimental validation (mediumpurple stars). **g,** The dose-response curves of experimentally synthesized compounds. Six molecules were successfully synthesized. The synthesis of 7a0983 was modified due to synthetic challenges, resulting in the production of 7a0983-100. **h,** The proposed binding modes of the successfully synthesized molecules with PI3Kα.

For the 96,240 innovative molecules, we conducted molecular docking using AutoDock Vina^41^. Although there is a known discrepancy between Vina scores and actual binding affinities, the Vina score remains an effective preliminary screening metric. Notably, 97.11% of these molecules exhibited a Vina score below -6.5, indicating that the novel molecules generated by ProLigGPT show promising potential for activity against PI3Kα (Fig.2c). We also calculated the maximum Tanimoto similarity on Morgan fingerprints^50^ between each of these innovative molecules and all known PI3Kα ligands from the training set. The average maximum similarity was 0.612 (SD = 0.170). Specifically, 84.30% of the generated molecules had a maximum similarity below 0.80, while 43.65% showed a maximum similarity below 0.60 (Fig.2d). These results suggest that the generated molecules are distinct and possess strong potential for effectively targeting PI3Kα, supporting the possibility of discovering PI3Kα ligands with novel scaffolds.

Our goal was to tackle the critical challenge of discovering ligands with novel scaffolds in drug design^51^. We preliminarily identified ligands with novel scaffolds as the generated molecules with completely distinct molecular fingerprints against the reference molecules, which consist of the known PI3Kα ligands and molecules from ligand repurposing (Fig.2a). The 1024-bit Morgan molecular fingerprints were used to cluster and compare the generated molecules with reference molecules (Methods), yielding a total of 846 clusters (Fig.2a, Extended Data Fig.5). Of these, 734 clusters containing reference molecules were excluded as they shared molecular fingerprints with the reference molecules, and therefore could not be considered novel scaffold ligands for further investigation (Fig.2e). The remaining 112 clusters became the focus of our exploration. These clusters were clearly distinct from the reference molecules, indicating they may contain previously unreported PI3Kα ligand scaffolds. To further confirm the stability and independence of these clusters, we performed the t-SNE^52^ analysis on these clusters and reference molecules (Extended Data Fig.6). Cluster 769 forms a tight cluster and exhibits clear separation from the reference molecules. It comprises 74 molecules that exhibited average maximum Tanimoto similarity values of 0.425 and 0.378 against all molecules appearing in the ProLigGPT training set and the kinase database^53,54^, respectively, confirming their notable novelty (Supplementary Materials Part 13). Further analysis identified a distinctive molecular scaffold, 1-(sulfonyl)-1*H*-benzimidazole, which has not been previously reported among ligands targeting PI3Kα or any other PI3K isoforms in the ChEMBL database^55^ (Fig.2f, Fig.S17). Subsequently, we selected seven molecules from Cluster 769 based on their molecular properties and docking results for experimental validation (Fig.2f, Extended Data Fig.3c, Supplementary Materials Part 14, Table S30). Six of these molecules were synthesized, and dose-response assays showed that all molecules inhibited PI3Kα effectively. Notably, four compounds, including 47546a, c89310, b42c5d, and dc081a (in-house ID, Fig.2g, Methods), demonstrated IC_50_ values below 10 μM, which indicates a significant potency in inhibiting PI3Kα. Compound c89310 is the most effective, with an IC_50_ value of 2.48 μM. Finally, to explore the inhibitor binding mechanism, we conducted molecular docking using AutoDock4^56^, generating 100 conformations for each synthesized molecule. By clustering these docking poses, followed by integrating predicted results with experimental activity data and expert insights, we inferred the binding mode between the synthesized molecules and PI3Kα (Fig.2h).

To sum up, ProLigGPT effectively avoided the issue of conservative outputs arising from training data constraints. The model successfully reproduced a wide variety of established ligand design strategies and generated inhibitors with novel scaffold structures beyond the patterns present in the training dataset. Through this capability, ProLigGPT explores uncharted chemical space, showcasing its unique strengths and broad potential for designing innovative drug candidates.

### ProLigGPT enables scaffold-based molecular design

In drug discovery, it is practically impossible to discover a new drug through brute-force exploration of chemical space, driving an increased demand for more rational approaches^57^. One such strategy involves starting with the core structures (scaffold) of known compounds to search for new molecules^58^. This approach, known as scaffold-based molecular design, focuses on designing derivative compounds that retain a particular scaffold as a substructure^59^. To address this demand, we enhanced ProLigGPT with the scaffold-based design mode through a prompt strategy, introducing a ligand prompt generation framework that allows users to specify a molecular scaffold represented by SMILES in addition to providing the protein sequence (Fig.1a, Opt2). By utilizing this framework, we can generate a library of derivative molecules by adding functional groups to the input molecular scaffold, offering a novel approach to finding molecules with desirable properties (Fig.3a, Supplementary Materials Parts 5, 11, and 12).

**Fig.3.**
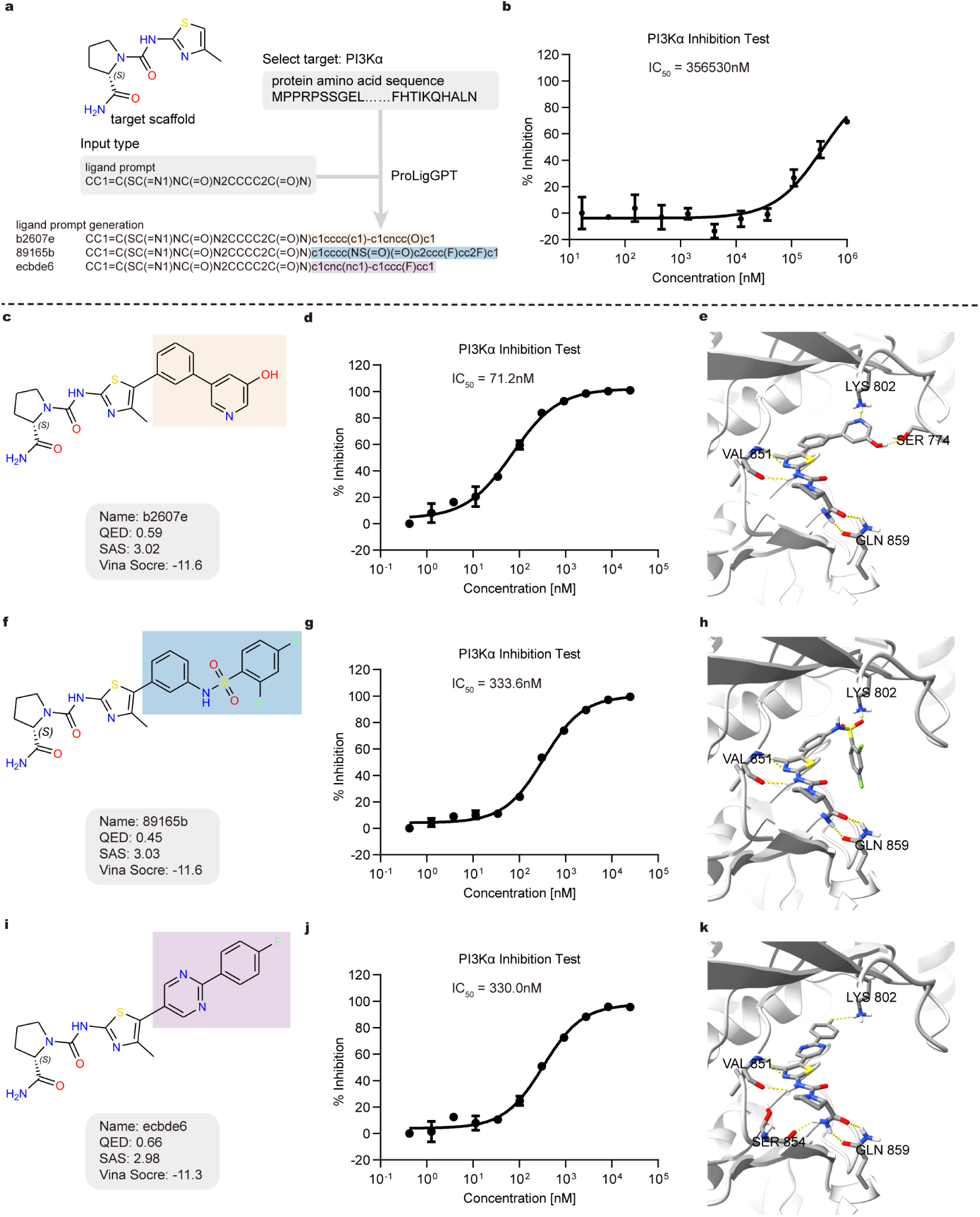
Using ligand prompt generation to decorate the seed scaffold into PI3K*α* ligands. **a**, Demonstration of ligand prompt generation. **b**, Dose-response curves of seed scaffold. **c,f,i,** Three example ligands generated using ligand prompt generation. **d,g,j,** Dose-response curves of the example ligands. **e,h,k,** The proposed binding mode of the example ligands with PI3Kα.

Similarly, using PI3Kα as a case, we generated derivative molecules based on the core scaffold of existing PI3Kα inhibitors and validated ProLigGPT’s effectiveness by wet-lab experiments. Alpelisib (BYL719, Extended Data Fig.7c) is a potent and selective PI3Kα inhibitor derived from the compound CHEMBL2071338 (Extended Data Fig.7f)^60,61^. Alpelisib preserves the key scaffold that builds upon attaching an (*S*)-pyrrolidine-2-carboxamide (L-Prolinamide) moiety via a urea linkage to 2-Amino-4-methylthiazole (PubChem CID: 74143)^61^. This scaffold confers the selective inhibiting α isoform of PI3K (Extended Data Fig.7i). Building on this, we provided ProLigGPT with the amino acid sequence of PI3Kα and the seed scaffold (key scaffold, Extended Data Fig.7i) as ligand prompt to generate 1000 derivative molecules (Fig.3a, Methods). Molecular docking of these derivative molecules with PI3Kα (PDB ID: 4jps) indicates that 96.39% of the molecules had a better Vina score than the seed scaffold, especially with 47.39% and 19.44% outperforming BYL719 and CHEMBL2071338, respectively (Extended Data Fig.7a). The molecular docking for BYL719 aligned well with its crystal structure (4jps, Extended Data Fig.7b), yielding an 0.57 Å r.m.s.d. (root mean squared deviation), particularly in the seed scaffold region, where a significant overlap was observed (Extended Data Fig.7d,e). Likewise, molecular docking analysis of CHEMBL2071338 and seed scaffold also showed good agreement with the crystal structure (Extended Data Fig.7g,h,j,k). The docking results suggest that molecules derived by adding functional groups to the seed scaffold may effectively interact with PI3Kα, while maintaining the interactions observed between the seed scaffold of BYL719 and PI3Kα.

Experimental validation is crucial to confirm the effectiveness of the scaffold-based molecular design using the ligand prompt generation strategy. We synthesized the seed scaffold (Extended Data Fig.6i) to use as a baseline, with dose-response assays yielding an IC_50_ of 356,530 nM (Fig.3b). To evaluate the effect of decorating the seed scaffold, we selected nine of the generated derivative molecules with better Vina scores than BYL719 and CHEMBL2071338 (Supplementary Materials Part 14, Table S31). Of these, eight molecules were synthesized and tested in dose-response assays, showing that five of them had IC_50_ values below 1μM (Extended Data Fig.8, Methods). For example, Compounds b2607e, 89165b, and ecbde6 (in-house ID) demonstrated desirable molecular properties and favorable molecular docking results, as well as effective inhibition of PI3Kα in experiments (Fig.3c–k). The docking results of these compounds also show similar interactions in the seed scaffold region to those observed with BYL719. Notably, b2607e had an IC_50_ value of 71.2 nM, and the functional groups generated by ProLigGPT within b2607e appear to interact with the SER774 site through hydrogen bonding, which represents a novel interaction not observed with CHEMBL2071338 and BYL719 (Fig.3c–e).

In addition to the ligand prompt generation strategy provided by ProLigGPT, derivative molecules based on the key scaffold of PI3Kα can also be generated using the R-group substitution from CADD. This alternative approach, however, relies on a carefully curated R-group replacement database. Here, we utilized the R-group replacement database developed by Kosuke Takeuchi et al.^62^ to generate 507 derivative molecules (Supplementary Materials Part 15). Molecular docking results demonstrated that these derivatives exhibited generally poor Vina scores, with only 0.79% surpassing BYL-719, and none exceeding CHEMBL2071338 (Fig.S19, Table S32). This performance is markedly inferior to the proportion of superior derivatives obtained through the ligand prompt strategy.

Altogether, the successful design of compounds with excellent Vina scores and significant biological activity clearly validates the effectiveness of ProLigGPT’s ligand prompt generation strategy. Starting from established scaffolds, ProLigGPT effectively extends known chemical spaces into adjacent uncharted chemical spaces, thus offering users a highly efficient and versatile platform for molecular design.

### ProLigGPT facilitates inhibitor discovery for structure-unresolved proteins with physiologically validated activity

Structural uncertainty remains one of the major barriers to structure-based drug design, particularly for proteins that are difficult to express, purify, and crystallize. To further demonstrate the unique capability of ProLigGPT in tackling such challenging targets, we selected Target T1. Target T1 is a validated therapeutic target, yet an experimentally determined 3D structure is unavailable for the target used in this study, rendering structure-based design impractical (Supplementary Materials Part 16). Moreover, its activity can be directly monitored by measuring analyte A1 levels in target-expressing cells, providing a physiologically relevant in-cell system for validating the functional activity of small molecules.

Using only the amino acid sequence of Target T1 (sequence code: S1), we applied the ProLigGPT *de novo* generation mode to design 10,000 candidate ligands without relying on structural information and molecular docking (Fig. 4a). After filtering by molecular properties (QED > 0.5, SA < 3.0), 4,223 molecules were retained (Fig. 4b). Exact search identified nine compounds in commercial libraries for validation in CL1 cells (Supplementary Materials Part 17), which endogenously express Target T1 and exhibit a measurable response in Assay M1 upon stimulation. Four ProLigGPT-designed compounds (C1, C2, C3, and C4) showed potent inhibition with IC50 values of 7.15, 1.64, 6.54, and 4.08 μM, respectively (Fig. 4c, e). Notably, three of these inhibitors reduced analyte A1 levels to basal levels comparable to unstimulated control cells, demonstrating full suppression of target activity. In contrast, R1, a known antagonist of Target T1 in other biological systems, inhibited Target T1 in the present system with an IC50 of 8.68 μM but failed to fully suppress the Assay M1 response even at 100 μM (Fig. 4d). To exclude cytotoxicity, direct interference with biosynthetic process B1 or nonspecific inhibition of enzyme E1, we performed Assays M2, M3 and M4, supporting the conclusion that the four ProLigGPT-generated compounds act as Target T1 inhibitors (Fig. 4f–i, Table S37).

**Fig.4.**
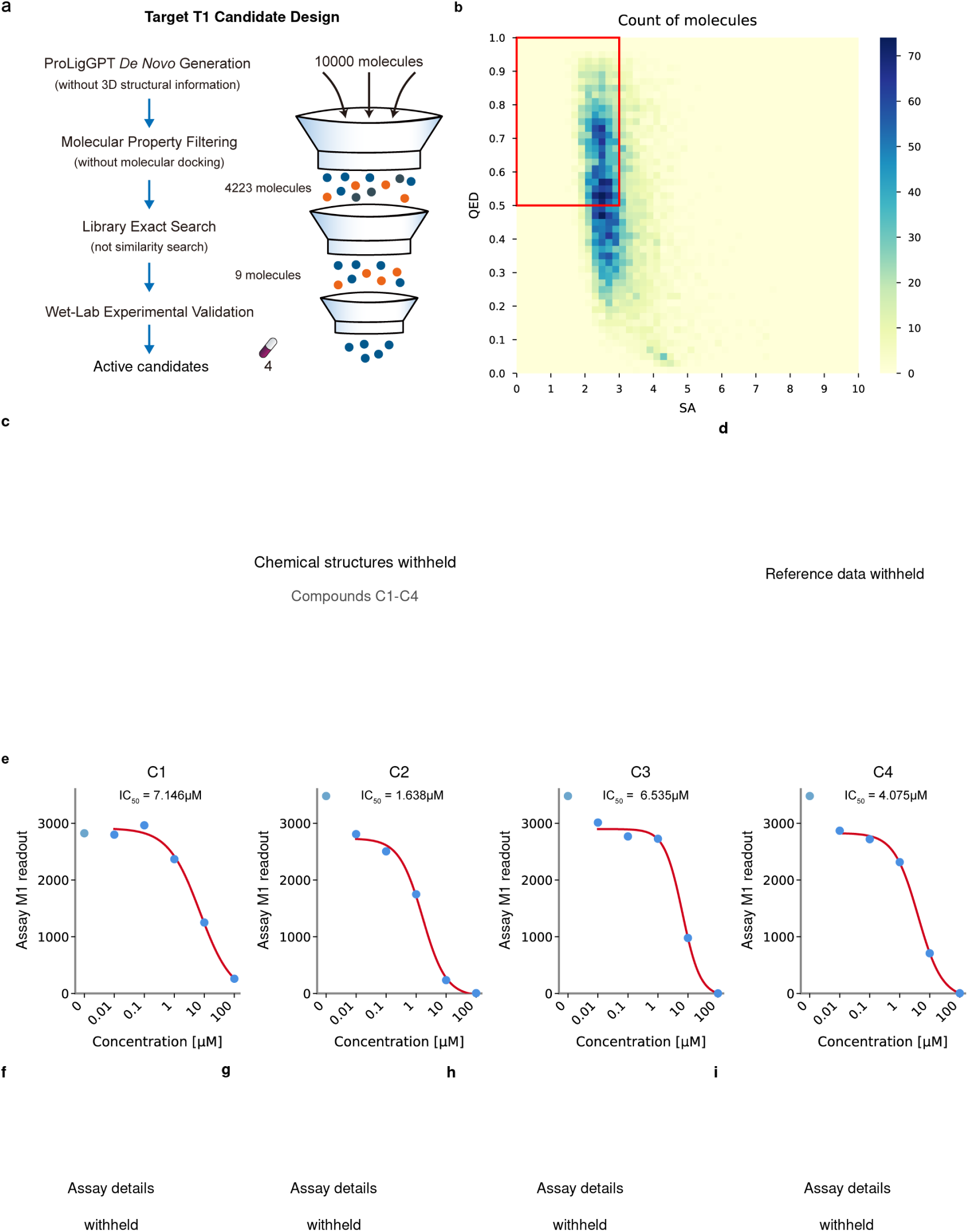
ProLigGPT discovers inhibitors for Target T1 without any structural information. **a,** ProLigGPT generated 10,000 candidate molecules using only the sequence of Target T1, without 3D structural information. Molecular-property filtering (QED > 0.5, SA < 3.0) retained 4,223 molecules; exact-match searches identified nine commercially available compounds for experimental testing. Four compounds showed concentration-dependent reductions in Assay M1. **b,** Distribution of generated molecules by QED and SA; the red box marks the filtering thresholds. **c,** Compound identities and chemical structures are withheld. d, The reference-compound comparison is withheld. **e,** Original dose-response observations for coded compounds C1, C2, C3 and C4 in Assay M1. The fitted IC50 values displayed in the figure are unchanged. The assay readout identity and original measurement units are withheld. **f–i,** Additional validation details are withheld. Target identity, biological context, assay procedures and the withheld panels are not available for independent assessment in this version. The retained observations do not independently establish target specificity or a mechanism of action.

These results demonstrate that ProLigGPT can design functionally active inhibitors for structurally unresolved targets solely from sequence information. The ability to discover effective inhibitors for Target T1, validated through in-cell assays, underscores the unique strength of ProLigGPT in generating truly functional small molecules without structural information. This case exemplifies ProLigGPT’s capacity to extend ligand discovery to structure-unresolved protein targets, thereby expanding the practical reach of sequence-based molecular design.

## Discussion

The development of ProLigGPT marks a significant advancement in molecular design. Without explicit pocket information, ProLigGPT outperforms structure-based models across targets with low sequence identity to proteins in its training set, achieving superior Vina scores while maintaining desirable drug properties such as QED and SA. TarPass benchmarking further shows that the model retains the largest overall candidate set after multi-stage screening across diverse target classes and generation paradigms, supporting robust generalization. This success stems from its sequence-to-sequence, decoder-only framework, which effectively leverages approximately 1.9 million experimentally validated protein-ligand sequence pairs. These extensive sequence data, which structural models cannot directly exploit, enable ProLigGPT to capture molecular bioactivity relationships solely through sequence-based learning, thereby overcoming limitations inherent to structure-based drug design. Additionally, by integrating *de novo* ligand generation and scaffold-based molecular design, ProLigGPT provides a powerful and accessible platform to explore previously underrepresented chemical spaces.

Unlike pocket-based design strategies, which typically focus solely on amino acids in direct proximity to the ligand-binding site, ProLigGPT utilizes full protein sequence information to generate ligands. Attention visualization, token composition analyses, shuffle experiments, and sensitivity assessments of the PI3Kα H1047R mutation collectively confirm this capability. This finding is consistent with the widely established principle that full-length protein sequences inherently encode structural information, first demonstrated by Anfinsen’s classical experiments^63^ and further validated by AlphaFold’s recent successes in predicting entire protein structures. Protein folding relies on cooperative interactions across the entire sequence; isolated residues near the pocket are insufficient to reliably determine stable conformations. Therefore, accurately capturing structural insights requires models to utilize complete sequences rather than limited local interactions.

Our emphasis on sequence information does not diminish the importance of pocket information but rather clarifies their underlying relationship. Pocket information is not an intrinsic protein attribute; it is a functional construct derived from integrating amino acid sequences, structural coordinates, functional annotations, and drug design knowledge. While sequences provide the foundational information, structural data (e.g., X-ray crystallography, cryo-EM, AlphaFold) provide spatial coordinates. Researchers then apply structural analysis, pharmacological data, and pocket detection algorithms to define clusters of residues likely involved in ligand binding as functional “pockets”^64^. Fundamentally, the intrinsic properties of a protein are determined entirely by its amino acid sequence, encoding all necessary information for folding and function^20^. Building upon this sequence-driven structural foundation, pocket information emerges as a specialized representation. It incorporates different prior knowledge, enabling the identification of distinct functional pockets within the same protein according to the specific ligand considered. However, pocket information does not introduce new structural or functional attributes beyond those stored within the amino acid sequence. Specifically, while ProLigGPT effectively responds to sequence variations, lacking explicit pocket-defined prior knowledge hinders its proactive control of ligand selectivity for specific mutations. To overcome this constraint, ProLigGPT incorporates a ligand prompt strategy that integrates prior interaction knowledge by retaining known selective scaffolds to enhance selectivity. Up to now, designing mutation-selective drugs has remained an open challenge, motivating our future work to focus on encoding pocket information into features usable by sequence models to improve precision and specificity.

Generative models often exhibit conservative behaviors when overly reliant on frequent patterns present in their training data, leading to mode collapse and text degeneration. Therefore, evaluating the innovation potential of models in such scenarios is crucial, and PI3Kα serves as an ideal case study. An innovative language model should generate new and experimentally validated active scaffolds that differ significantly from known ligands, even after fully learning existing ligand data for target proteins. This capability clearly shows the model’s comprehension and creativity. ProLigGPT achieves this by producing a cluster of experimentally validated compounds that share a novel PI3Kα inhibitor scaffold, and these compounds are collectively distinct from all molecules present in both its training data and existing kinase inhibitor libraries. These results differ from situations in which obtaining a single inhibitor with a novel scaffold might be attributed merely to chance, clearly illustrating ProLigGPT’s ability to propose and utilize novel scaffolds for inhibitor design. This ability effectively addresses one of the fundamental challenges in drug discovery: generating innovative chemical structures^51^.

Unlike the encoder-decoder architecture, ProLigGPT leverages ligand prompts for conditional control, enabling flexible optimization strategies. Compared to R-group substitution, which relies on predefined fragment databases, lacks protein specificity, and offers limited structural combinations, ProLigGPT’s ligand prompt strategy utilizes only protein sequences and core scaffolds, enabling it to propose molecular derivatives and novel interactions that significantly broaden chemical exploration. In the PI3Kα example, ProLigGPT automatically modified the key scaffold of CHEMBL2071338 by rationally adding functional groups, producing compound b2607e (IC₅₀ = 71.2 nM). This compound exhibited a 5000-fold increase in scaffold activity, significantly surpassing CHEMBL2071338, and demonstrated potential novel hydrogen bond interactions with residue SER774. Although b2607e has not yet surpassed BYL719, a compound extensively refined through interdisciplinary collaboration involving medicinal chemistry, structural biology, and expert-driven optimization, ProLigGPT rapidly produced b2607e by decorating the key scaffold within a short timeframe using fully automated sequence-based methods. This underscores ProLigGPT’s notable capacity to accelerate early-stage drug discovery.

Finally, we experimentally validated ProLigGPT’s generalization to Target T1 from the species studied, a previously unseen GPCR with no reported crystal structure or known inhibitors. GPCRs remain among the most challenging targets in structural biology because low expression limits protein yield while membrane extraction compromises receptor stability, with conformational flexibility making it difficult to stabilize the receptor in a single conformation^65^. Among 401 human non-olfactory GPCRs, only 112 have experimental structures; of these, 36 have been resolved in both active and inactive states^66^. Structure-dependent workflows requiring receptor models, predefined pockets, or molecular docking are difficult to extend to non-human receptors, where experimental structure coverage is even lower. This constraint is compounded by a broader pharmacological gap in animal-model research, where protein sequence data, cellular models with endogenous target expression, large-scale phenotyping datasets, and established disease models are widely available, yet species-matched small-molecule tools for endogenous targets remain scarce. Unlike gene knockout approaches that reveal the consequences of gene loss, small-molecule intervention addresses a distinct biological question by testing how an established phenotype responds to acute, reversible, dose-dependent changes in endogenous protein activity. Because protein sequences are widely available across animal species, generating candidate molecules directly from sequence offers a scalable route to species-matched pharmacological tools for previously unseen targets. To test this approach, using the Target T1 sequence as the sole target input, ProLigGPT generated candidates that were subsequently filtered by molecular properties. Experimental testing of commercially available exact matches yielded four micromolar inhibitors, demonstrating direct molecular design for an endogenous animal protein. As far as we know, this is the first experimental demonstration that small-molecule inhibitors can be discovered directly from the protein sequence of a GPCR lacking an experimental structure. Neither molecular generation nor candidate selection relied on structural modeling, pocket definition, or molecular docking. This result establishes the feasibility of the “sequence → ligand” end-to-end paradigm, which differs from multi-model integrated pipelines such as “sequence → structure → pocket → ligand”, providing a more direct route for molecular design when structural information is unavailable. The commercial matches also show that ProLigGPT samples synthetically accessible and immediately testable chemical space. Experimental testing of these exact matches yielded inhibitors, illustrating how ProLigGPT can awaken molecules of previously unknown function within chemical libraries, reveal their potential targets, and transform them into pharmacologically meaningful compounds. Overall, these results demonstrate that language models can design functional ligands directly from target protein sequences, entirely free from structural information. In conclusion, computational evaluations and experimental validations collectively demonstrate the model’s intrinsic ability to support cross-domain, multi-task applications, effectively overcoming structural constraints encountered in molecular design. Although the decoder-only architecture inherently permits the incorporation of quantitative bioactivity labels for affinity prediction, ProLigGPT currently ensures predictive reliability through post-generation evaluation: proteins with available structural data undergo molecular docking validation, whereas those lacking structural data utilize sequence-based predictive models such as TransformerCPI2.0^67^. Moving forward, based on the demonstrated performance and flexibility of ProLigGPT, the framework aims to expand to a broader spectrum of molecular entities, including nucleic acids^68^, peptides^69^, antibodies^70^, and PROTACs^71^, concurrently incorporating strategies such as molecular linker design^72^, scaffold hopping^73^, and affinity prediction to further enhance its capabilities. As language models continue to evolve, their integration with experimental workflows is expected to accelerate therapeutic drug discovery and development, providing innovative solutions to precision medicine and global health challenges.

## Conclusion

In this study, we developed ProLigGPT, a novel sequence-based molecular design model that leverages the full protein sequence to generate bioactive molecules without relying on structural data. By employing a seq2seq autoregressive architecture based on GPT-2^27^, ProLigGPT excels in both *de novo* and scaffold-based drug design. Our results demonstrate that ProLigGPT outperforms structure-based drug design models in several key metrics, while offering greater flexibility in exploring chemical space and generating novel molecules with desirable drug-like properties. Interpretability analysis indicates that ProLigGPT uses the full protein sequence and exhibits sensitivity to point mutation, unlike pocket-based models that depend only on residues near the binding site. Through experimental validations, ProLigGPT successfully designed inhibitors sharing a novel scaffold and inhibitors derived from a known scaffold for PI3Kα, expanding the chemical space beyond existing compounds. Additionally, ProLigGPT generated promising compounds with strong activity against Target T1, a structure-unresolved membrane protein without known inhibitors, highlighting ProLigGPT’s ability to design inhibitors for even the most challenging targets free from structural constraints. Overall, ProLigGPT demonstrates success in utilizing protein sequences for molecular design, marking a significant step toward the future of sequence-based ligand discovery and paving the way for more efficient drug candidate generation.

## Methods

### Data source

The data used in our study were sourced from the Hugging Face Dataset Hub (https://huggingface.co/datasets), specifically the ‘zpn/zinc20’ (https://huggingface.co/datasets/zpn/zinc20) and ‘jglaser/binding_affinity’ (https://huggingface.co/datasets/jglaser/binding_affinity) datasets. Both datasets were loaded and processed using the Hugging Face Datasets library^74^ (https://github.com/huggingface/datasets). The ‘zpn/zinc20’ dataset comprises approximately one billion commercially available compounds from the ZINC database^75^, represented as SMILES and SELFIES^76^ strings. The ‘jglaser/binding_affinity’ dataset contains 1.9 million unique protein-ligand pairs with experimentally determined binding affinities, aggregated from databases including BindingDB^77^, PDBbind-cn^78^, BioLIP^79^, and BindingMOAD^80^. We exclusively chose SMILES over SELFIES due to its broader compatibility with chemical toolkits and demonstrated advantages in recent molecular generation benchmarks^81^.

### ProLigGPT tokenizer

Given the significant differences between molecular design data and natural language data, training a tokenizer from scratch is more suitable for our research than using existing tokenizers like GPT-2. Consequently, the ProLigGPT tokenizer was developed from scratch using the ‘jglaser/binding_affinity’ dataset. We created distinct tokenizers for proteins and ligands separately, and after merging, deduplication, and additional preprocessing steps, we built the ProLigGPT tokenizer. This approach enables ProLigGPT to effectively process both biological sequences from protein amino acid sequences and chemical sequences represented by ligand SMILES, thus bridging the gap between biological and chemical sequence domains.

#### Tokenization of proteins

We initially trained a tokenizer using the BPE algorithm on the protein-ligand pair data derived from the ‘jglaser/binding_affinity’ dataset, formatted as ‘<P>’ + protein amino acid sequence + ‘<L>’ + ligand SMILES, with the vocabulary size set to 50,000 inspired by GPT-2 and ProtGPT2^20^. After training, we observed that the resulting vocabulary predominantly consisted of protein-related fragments, reflecting the higher complexity of protein amino acid sequences compared to ligand SMILES sequences. Hence, despite the initial inclusion of ligand data, we refer to this tokenizer as the protein tokenizer for simplicity.

#### Tokenization of ligands

Since ProLigGPT is designed specifically to generate ligand SMILES, we separately trained a ligand tokenizer using the BPE algorithm on the ligand portion of the ‘jglaser/binding_affinity’ dataset, explicitly aiming to enhance ligand vocabulary representation. In line with the protein tokenizer, the ligand tokenizer vocabulary size was initially set to 50,000; however, training produced only 3,560 unique tokens (excluding special tokens). The difference in vocabulary size between the ligand tokenizer and protein tokenizer aligns with the distinct complexities of ligand SMILES and protein sequences.

#### Building the ProLigGPT tokenizer

To enable the ProLigGPT tokenizer to handle both molecular SMILES sequences and protein sequences, we merged the ligand tokenizer with the protein tokenizer. Initially, overlaps existed between the vocabularies of the two tokenizers: the protein tokenizer contained 50,000 vocabulary items (excluding the special token ‘<|endoftext|>’ and the 256 initial characters, leaving 49,743 unique tokens), and the ligand tokenizer contained 3,560 vocabulary items (excluding the 256 initial characters, leaving 3,304 unique tokens). Specifically, there were 223 overlapping vocabulary items between the 49,743 unique tokens of the protein tokenizer and the 3,304 unique tokens of the ligand tokenizer. To resolve potential conflicts, we removed these overlapping tokens along with their corresponding merge rules from the protein tokenizer, leaving 49,520 unique tokens. We then added the 3,304 unique ligand tokenizer tokens and their associated merge rules, resulting in a combined vocabulary of 52,824 tokens. Finally, we incorporated the initial 256 characters from the BPE algorithm and three special tokens (’[PAD]’, ‘<|startoftext|>’, and ‘<|endoftext|>’), finalizing the ProLigGPT tokenizer with a total vocabulary size of 53,083. The ProLigGPT tokenizer supports encoding, decoding, and tokenization functionalities for ProLigGPT model inputs and outputs. For additional details, please refer to the Supplementary Materials.

### ProLigGPT training datasets

In this study, we utilized two datasets to train the ProLigGPT model: the Molecule Text dataset generated by ‘zpn/zinc20’ and the Protein-Ligand Pair Text dataset generated by ‘jglaser/binding_affinity’. In the training process, one epoch is divided into two stages: First, we used the Molecule Text dataset to train ProLigGPT, enhancing its ability to understand and generate chemical representations of ligands. In the second stage, we continued training with the Protein-Ligand Pair Text dataset. This further improves the model’s ability to generate ligands, enhances its understanding of protein sequences, and helps capture potential activity relationships between proteins and ligands. This approach facilitates the generation of small molecules that bind to target proteins.

#### Molecule Text dataset

We extracted SMILES sequences from the ‘zpn/zinc20’ dataset. Each SMILES sequence was then converted into molecule text data in the format ‘<|startoftext|>’ + ‘<L>’ + SMILES sequence + ‘<|endoftext|>’. Subsequently, we used the ProLigGPT tokenizer to encode the text data. To ensure the randomness and representativeness of the data, we shuffled the entire dataset (1,006,650,595 molecule text entries). After shuffling, we allocated the final 10,067 entries (0.001%) as the validation set, and for each epoch, we randomly sampled 100,666 entries (0.01%) from the remaining data as the training set.

#### Protein-Ligand Pair Text dataset

We extracted protein-ligand pairs from the ‘jglaser/binding_affinity’ dataset and converted each pair into text data in the format ‘<|startoftext|>’ + ‘<P>’ + protein amino acid sequence + ‘<L>’ + ligand SMILES + ‘<|endoftext|>’. Subsequently, we encoded this data using the ProLigGPT tokenizer. To ensure randomness, we shuffled the entire dataset. We split the entire dataset into a training set and a validation set at a 9:1 ratio.

### ProLigGPT model training details

To train the ProLigGPT model for ligand discovery, we selected the GPT2LMHeadModel from Hugging Face Transformers library^82^ as our base model. The superior ability of GPT2LMHeadModel to capture complex language patterns is advantageous for representing chemical and biological sequences. GPT2LMHeadModel extends GPT2Model by adding a language modeling head that maps hidden state vectors to the vocabulary size. This addition enables the model to predict the next token in a sequence, with training guided by the language modeling loss function designed for next-token prediction. We used the Hugging Face Transformers and Datasets libraries, which provide powerful NLP tools and integrate seamlessly with PyTorch, enhancing our training efficiency.

Because the sequence information of small molecules and proteins is fundamentally different from natural language text, we chose to train the model from scratch rather than fine-tune a pre-trained language model. Training from scratch allows the model to better capture the unique features inherent in chemical and biological sequences. The model was configured with a maximum sequence length of 1,024 tokens, an embedding and hidden state dimensionality of 768, 12 hidden layers, and 12 attention heads per layer. The detailed architecture of the ProLigGPT model is provided in the Supplementary Materials. We employed a language modeling loss function designed for next-token prediction to train the model effectively for sequence generation tasks. The training was conducted on an NVIDIA RTX 4090 GPU. We used the AdamW optimizer with a learning rate of 5e-4, an epsilon value of 1e-8, and a warm-up period of 100 steps. Additionally, we implemented a linear learning rate scheduler with a warm-up using the get_linear_schedule_with_warmup function to adjust the learning rate dynamically during training.

The training was divided into two stages. The first stage involved training the model using the Molecular Text dataset to teach it how to accurately use SMILES representations of molecules. The second stage used the Protein-Ligand Pair Text dataset to instruct the model on generating compounds for target proteins. This requires the model to understand protein amino acid sequences and generate compounds that interact with them. For batch processing during training, we standardized the length of the data using the ProLigGPT tokenizer. We decided to truncate and pad the data to a length of 768, which reduces computational load while covering 98.62% of the Protein-Ligand Pair Text dataset (Supplementary Materials Part 3).

We set the training for 5 epochs with a batch size of 4, while splitting the Protein-Ligand Pair Text dataset into training and validation sets in a 9:1 ratio. The training process took approximately one week. After five training cycles, the model’s loss on the validation set dropped to 0.04, an encouraging result indicating that ProLigGPT can effectively understand and generate compounds for specific proteins.

### ProLigGPT generation

During the generation process, the model selects the next word from its predicted probability distribution according to the set sampling strategy. These strategies involve multiple parameters, including top_k, top_p, and temperature. In ProLigGPT, the default value for top_k is 9, meaning the model only chooses from the top 9 most probable words at each step of generation. The default value for top_p is set to 0.9, indicating that the model will select words until the cumulative probability reaches 0.9. The default setting for temperature is 1.0. Reducing the temperature increases the likelihood of choosing higher-probability words while decreasing the likelihood of lower-probability ones, making the distribution steeper and enhancing generation accuracy. Increasing the temperature flattens the distribution, enhancing creativity. Combining these strategies allows the model to dynamically adjust to meet the needs of different scenarios.

Additionally, other parameters need to be configured during the generation process. The device parameter is set to ‘cuda’ by default, as GPUs significantly enhance running efficiency compared to CPUs. The default batch_size is 16. It should be adjusted based on GPU performance to avoid exceeding memory limits or experiencing slowdowns due to shared memory. The number parameter specifies how many molecules will be returned. The min_atoms and max_atoms parameters control the minimum and maximum number of non-hydrogen atoms in the generated molecules, with default values set to None and 35, respectively. However, these two parameters are not effective in the ligand prompt generation mode.

Adapting parameters to align with specific research needs is essential for achieving diverse objectives. For example, we used default parameters during model evaluation; when exploring novel scaffold ligands for PI3Kα, we adjusted parameters to enhance creativity. Detailed command line instructions for specific parameter choices are provided in the Supplementary Materials.

To avoid conflicts that might arise from numerical naming, we adopted a unique identifier system using the SHA-1 hash value of the generated molecule’s SMILES string. This approach effectively resolves issues with naming conflicts (Supplementary Materials Part 11).

#### ProLigGPT Colab Notebook

To enable more people to conveniently use ProLigGPT, we have deployed an interactive notebook titled “ProLigGPT_Colab.ipynb” publicly accessible on Google Colab (https://colab.research.google.com/drive/1lxDpEGz9498tGvEi7nSgryXI4haMKMhc). Despite the installation process of ProLigGPT being fairly straightforward, deployment on Google Colab allows users unfamiliar with the installation process to use ProLigGPT easily. Moreover, it facilitates the utilization of cloud-based GPU resources, significantly improving operational efficiency, especially for users without GPU resources. Within ProLigGPT_Colab, we provide five main steps and a check step to guide users in smoothly operating ProLigGPT. The check step, namely Step 0: Check GPU, is the initial process. This notebook requires running in GPU mode, and users can execute the nvidia-smi command to verify if the GPU has been correctly loaded. The five main steps are as follows:

Step 1. Clone Repository: Clone the source code of ProLigGPT from GitHub and set the current working directory to the ProLigGPT directory.

Step 2. Build Environment: Install all the dependent libraries required to run ProLigGPT.

Step 3. Run ProLigGPT: Users only need to complete the relevant settings during runtime as prompted, and they can successfully run ProLigGPT with one click.

Step 4. Save Results: Users can choose to download the necessary results folders to their local computers.

Step 5. Visualize and Post-process on Your Local Computer: We provide some suggestions to guide users in visualizing and post-processing results on their local computers.

These five steps are simple and user-friendly; users only need to follow the guide on Google Colab to easily complete them. By deploying ProLigGPT on Google Colab, we provided a convenient and easy-to-use platform, enabling more people to apply ProLigGPT seamlessly in drug discovery research.

#### ProLigGPT Command Line Interface

ProLigGPT can also be deployed locally. This option is ideal for users who possess substantial computational power and wish to run the model or conduct large-scale calculations on their own machines. Such users can fully leverage the advantages of their local computing resources to achieve higher computational efficiency and more flexible scheduling strategies. Even those with a moderate-performance GPU can use their local devices to generate bioactive molecules. ProLigGPT supports running under both Windows and Linux, ensuring compatibility across operating systems. To run ProLigGPT locally, please visit our GitHub repository at https://github.com/LIYUESEN/ProLigGPT, download the ProLigGPT code, complete the deployment, and follow the instructions in the README document or the guidance from running ‘python proliggpt.py -h’.

### ProLigGPT required post-processing

The required post-processing is primarily conducted through the Open Babel library. It involves two key stages: Converting ligand SMILES to SDF and Energy minimization. This process aims to ensure that the generated molecular structures are chemically valid and stable.

#### Convert ligand SMILES to SDF

In this study, we primarily use two chemical structure representation formats: SMILES and SDF. SMILES is a method for representing chemical structures using simple ASCII strings. It employs atomic symbols, bond connections, ring structures, and parentheses to depict molecular structures in a concise and readable manner. The fundamental principle of SMILES is to describe the molecule’s topology through a linear representation of atoms and bonds. SDF (Structure-Data File) is a format used to store and transmit chemical structure data, providing detailed three-dimensional information, including atomic coordinates, bond connections, and other molecular details. In our research, the SDF format is used to store the structures of generated molecules for subsequent analysis and experimental validation.

During ProLigGPT generation, the output of the ProLigGPT model is decoded and converted by the ProLigGPT tokenizer into the molecular SMILES representation. For structural analysis and experimental validation, the required post-processing step includes converting the generated SMILES into SDF format. This conversion is typically carried out using open-source cheminformatics libraries such as Open Babel. Converting SMILES to SDF format allows us to thoroughly explore and visualize the three-dimensional structural features of the molecules. This process also ensures their chemical validity and stability.

#### Energy minimization

The generated molecules, after being converted from SMILES to SDF format, cannot be directly used for subsequent analysis. This initial conversion only ensures the molecular atomic types and bond connections, representing the topological structure, without considering energy aspects. The thermodynamically stable structure of a molecule is typically represented by its lowest energy state, as molecules naturally exist in their most stable form. The energy minimization process simulates the most stable configuration of the molecule in its natural state. Therefore, in molecular docking experiments, the small molecule structures also need to undergo energy minimization. Consequently, in the Required Post-processing steps, the Open Babel library (specifically, the openbabel module from Open Babel v3.0.0, providing access to almost all of the Open Babel interfaces via Python) is used to perform energy minimization on the generated molecules (conformational optimization only, without altering the molecular state). The MMFF94 force field^83^ is employed, and 10,000 steps of the steepest descent method are performed. These steps ensure the chemical validity and stability of the molecular structures we generate.

### ProLigGPT attention visualization with BertViz

To visualize ProLigGPT’s attention and enhance the interpretability of generated molecules, we utilized BertViz^84^, an interactive tool designed to visualize attention in Transformer-based models. BertViz enables intuitive exploration of attention distributions, providing clear insights into the model’s decision-making processes.

ProLigGPT employs a 12-layer decoder architecture, with each layer equipped with attention to facilitate sequence-to-sequence modeling. Specifically, the first layer’s attention directly operates on input token embeddings, and its attention weights explicitly illustrate how the model initially focuses on original input tokens. Attention in subsequent layers acts upon hidden states generated by preceding layers, which have undergone several complex transformations integrating contextual, positional, higher-level functional information, and long-range dependencies. Consequently, attention weights at higher layers reflect complex semantic combinations rather than the original meanings of input tokens.

We developed a Colab notebook integrating BertViz’s head_view and model_view functionalities (https://colab.research.google.com/drive/1HUioWgOmVxmoPsl-9qbyFlWajCdTHa) for intuitive visualization of attention distributions. This notebook explicitly visualizes ProLigGPT’s first-layer attention, illustrating clearly how the model directly leverages input token information. Additionally, it provides comprehensive visualizations for all 12 attention layers, enabling users to deeply analyze how information is progressively processed and propagated within the model. By inputting the target protein sequence (e.g., PI3Kα, UniProt ID: P42336) and a generated molecule represented in SMILES format (e.g., c89310) into our Colab notebook, users can intuitively explore the model’s generation process via an interactive visualization interface.

### t-SNE visualization

The generated molecules are converted from SMILES representations to 1024-dimensional Morgan fingerprints using the Open Babel library. Subsequently, the t-SNE technique from the scikit-learn library is used for data dimensionality reduction and visualization, with the settings: n_components=2, perplexity=30, n_iter=100000, and random_state=42.

### Louvain method for community detection

To identify community structures within the Morgan fingerprints of generated molecules, the nearest neighbors are computed first using the NearestNeighbors function from the scikit-learn library^85^ with k=10 and a Minkowski metric. Then a graph is constructed with the networkx library^86^, where nodes represent individual molecules and edges connect nearest neighbors. The edge weights are set inversely proportional to the distances. Finally, the Louvain method^87^ is applied for community detection using the community_louvain library, optimizing modularity with settings: resolution=20 and random_state=42.

### Chemical synthesis and analytics

The synthesis of the compounds was carried out by Pharmaron Qingdao Co., Ltd. (Address: Building C, No. 7 Fenglong Road, High-tech Zone, 266114 Qingdao, Shandong, China). The company supplied detailed analytical data, including 1H NMR, 13C NMR, and LCMS, to verify the identity and purity of the synthesized compounds. For a comprehensive overview of the chemical synthesis and compound analytics, please refer to Chemical Synthesis and Dose-Response Analysis.

### Dose-response assay and IC50 determination for PI3K*α*

The dose-response assay and IC_50_ determination were performed by Pharmaron Beijing Co., Ltd. (Address: 6 TaiHe Road, BDA, Beijing, 100176, P.R. China) following standard protocols. To evaluate the IC_50_ of the synthesized compounds against PI3Kα, the compounds were tested across 10 gradient concentrations. IC_50_ values were calculated by fitting the % Inhibition and the log of compound concentrations to a nonlinear regression (dose response – variable slope) using GraphPad Prism 8. Details on initial concentrations, dilution factors, as well as materials and reagents are provided in Chemical Synthesis and Dose-Response Analysis.

### Experimental information for Target T1

The target identity, biological system, compound identities, assay readouts, measurement units, reagents and experimental procedures for the Target T1 case study are withheld in this version pending intellectual-property protection.

### Dose-response analysis

For the dose-response observations retained in Fig. 4e, IC50 values were estimated by fitting the assay readout against compound concentration using four-parameter variable-slope nonlinear regression.

## Data availability

The data used in our study was sourced from the Hugging Face Dataset Hub. The ligand SMILES data was sourced from the ‘zpn/zinc20’ database: https://huggingface.co/datasets/zpn/zinc20. Unique pairs of protein sequences and ligand SMILES data were obtained from the ‘jglaser/binding_affinity’ database: https://huggingface.co/datasets/jglaser/binding_affinity. Data availability is under review. The identity, compound structures, and detailed experimental methods of Target T1 are withheld.

## Code availability

Our code and checkpoints employed in this research are openly accessible to the public. They are hosted on two platforms, GitHub (https://github.com/LIYUESEN/ProLigGPT) and Hugging Face (https://huggingface.co/liyuesen/proliggpt), to maximize accessibility and convenience for future use and replication. An interactive and easy-to-use Google Colab notebook for running ProLigGPT is publicly available at https://colab.research.google.com/drive/1lxDpEGz9498tGvEi7nSgryXI4haMKMhc. An interactive visualization of ProLigGPT implemented in a Google Colab notebook is available at https://colab.research.google.com/drive/1HUioWgOmVxmoPsl-9qbyFlWajCdTHa.

## Acknowledgments

We thank Dr. Nan Sheng, Dr. Yuyang Zhang, Changcheng Wang, Xuesong Han, Panwei Yu, and Jiale Wang for their insightful discussions and feedback. We also thank Pharmaron Qingdao Co., Ltd. for chemical synthesis support, in addition to Pharmaron Beijing Co., Ltd. for dose-response analysis support.

## Author contributions

Y.S.L. proposed the original idea and implemented the main code. Y.S.L., X.S. deployed ProLigGPT on Colab. X.W. improved the code. Y.S.L. collected and processed the data. Y.S.L., X.W., and X.S. tested the code. Y.S.L. designed the computational experiments. Y.S.L., X.S., and X.W. performed computational experiments. Y.S.L., C.G., X.S., and X.W. analyzed the results of computational experiments. Y.P.L., Y.S.L., X.T., and W.M. designed, supervised, and analyzed the wet-lab experiments for PI3Kα. P.X., K.X., H.C., H.F., J.M., and Y.S.L. designed, performed, and analyzed the wet-lab experiments for Target T1. Y.S.L., C.G., and X.S. wrote the initial draft of the manuscript. Y.S.L., C.G., and X.W. created the figures for the manuscript. Y.X., Y.S.L. wrote the main manuscript. Y.X. led the manuscript revision and the responses to reviewers, with contributions from J.M., Y.S.L., F.W., J.L., Y.T.L., H.Z., and Y.W. All authors contributed to the manuscript. Y.X., Y.P.L., J.M., and S.H. provided advice and guidance on the project. S.H. contributed to the essential financial support and led the project.

## Competing interests

S.H., Y.X., Y.S.L., C.G., X.S., and X.W. have been granted a Chinese patent (Patent No.: ZL 2024 1 0754023.5) in the name of the First Affiliated Hospital of Xi’an Jiaotong University, relating to the autoregressive model-based drug design method. S.H., Y.X., Y.S.L., C.G., Y.P.L., X.S., and X.W. have been granted a U.S. patent (Patent No.: US 12,437,847 B1) in the name of the First Affiliated Hospital of Xi’an Jiaotong University, relating to the autoregressive model-based drug design method. S.H., Y.P.L., Y.X., Y.S.L., and C.G. have filed a U.S. patent application in the name of the First Affiliated Hospital of Xi’an Jiaotong University, relating to a series of PI3Kα inhibitors generated based on the ProLigGPT model. The authors declare that they are preparing a patent application related to the Target T1 inhibitors generated based on the ProLigGPT model.

## Supplementary information

Supplementary File 1: Supplementary Materials

Supplementary File 2: Chemical Synthesis and Dose-Response Analysis

## Extended Figures

**Extended Data Fig.1.**
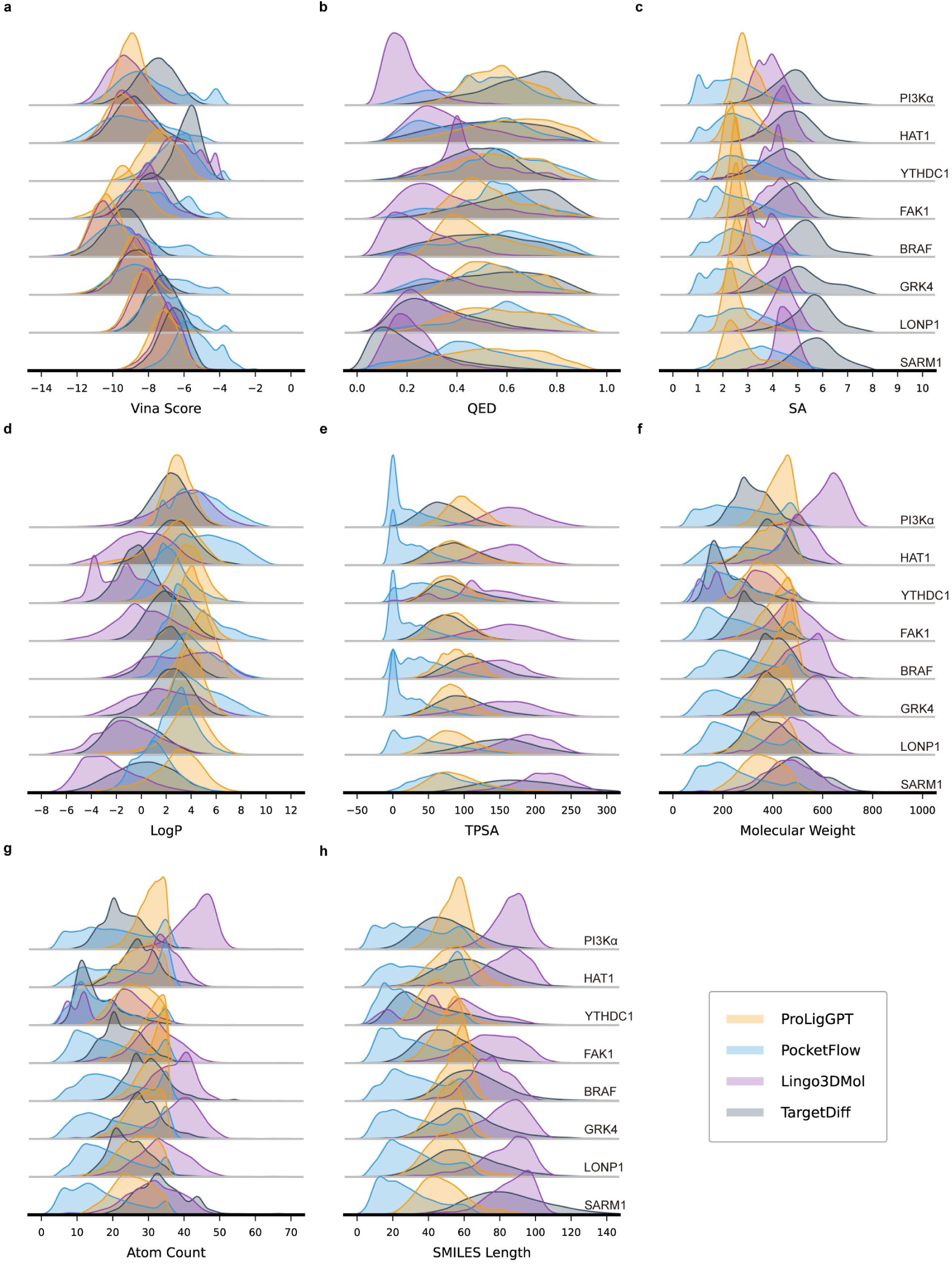
The distributions of molecular properties and metrics across four models for eight target proteins. **a–h,** The distributions of eight molecular properties and metrics for molecules generated by four distinct drug design methods (ProLigGPT, PocketFlow, Lingo3DMol, and TargetDiff), using default settings for eight protein targets (PI3Kα, HAT1, YTHDC1, FAK1, BRAF, GRK4, LONP1, and SARM1), are presented, including Vina score (**a**), QED (**b**), SA (**c**), logP (logarithm of partition coefficient, **d**), TPSA (topological polar surface area, **e**), molecular weight (**f**), atom count (**g**), and SMILES length (**h**).

**Extended Data Fig.2.**
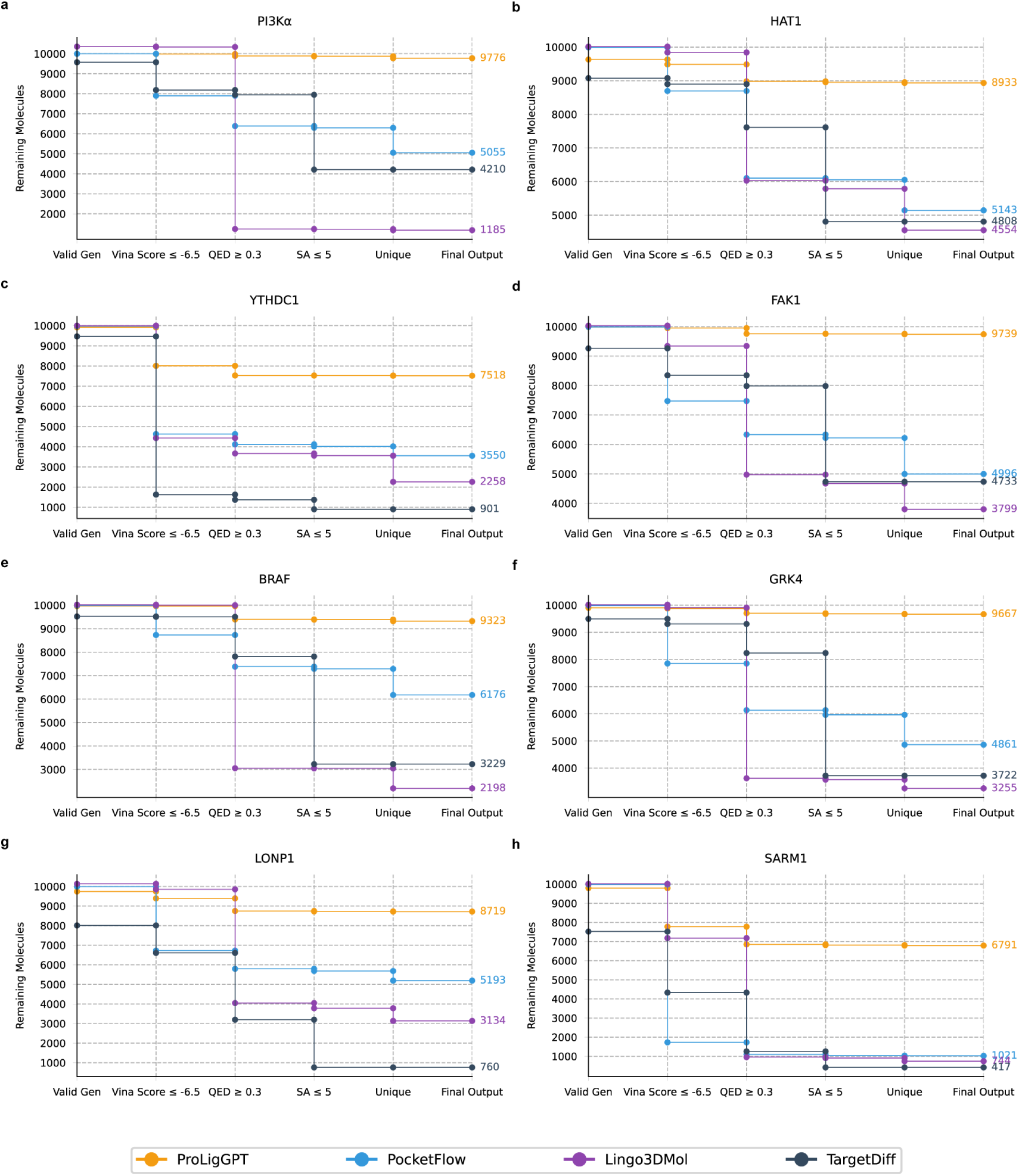
ProLigGPT outperforms structure-based models in generating drug-like targeted molecules. **a–h,** ProLigGPT was compared with three structure-based models (PocketFlow, Lingo3Dmol, TargetDiff) by individually generating 10,000 molecules for eight different types of target proteins, including PI3Kα (**a**), HAT1 (**b**), YTHDC1 (**c**), FAK1 (**d**), BRAF (**e**), GRK4 (**f**), LONP1 (**g**), and SARM1 (**h**). The generated molecules were then going through sequential filtering to evaluate their potential binding affinities and drug-like properties, including Vina score ≤ -6.5, QED ≥ 0.3, SA ≤ 5, and Uniqueness. After filtering, the model with more molecules retained demonstrates better performance in generating drug-like targeted molecules.

**Extended Data Fig.3.**
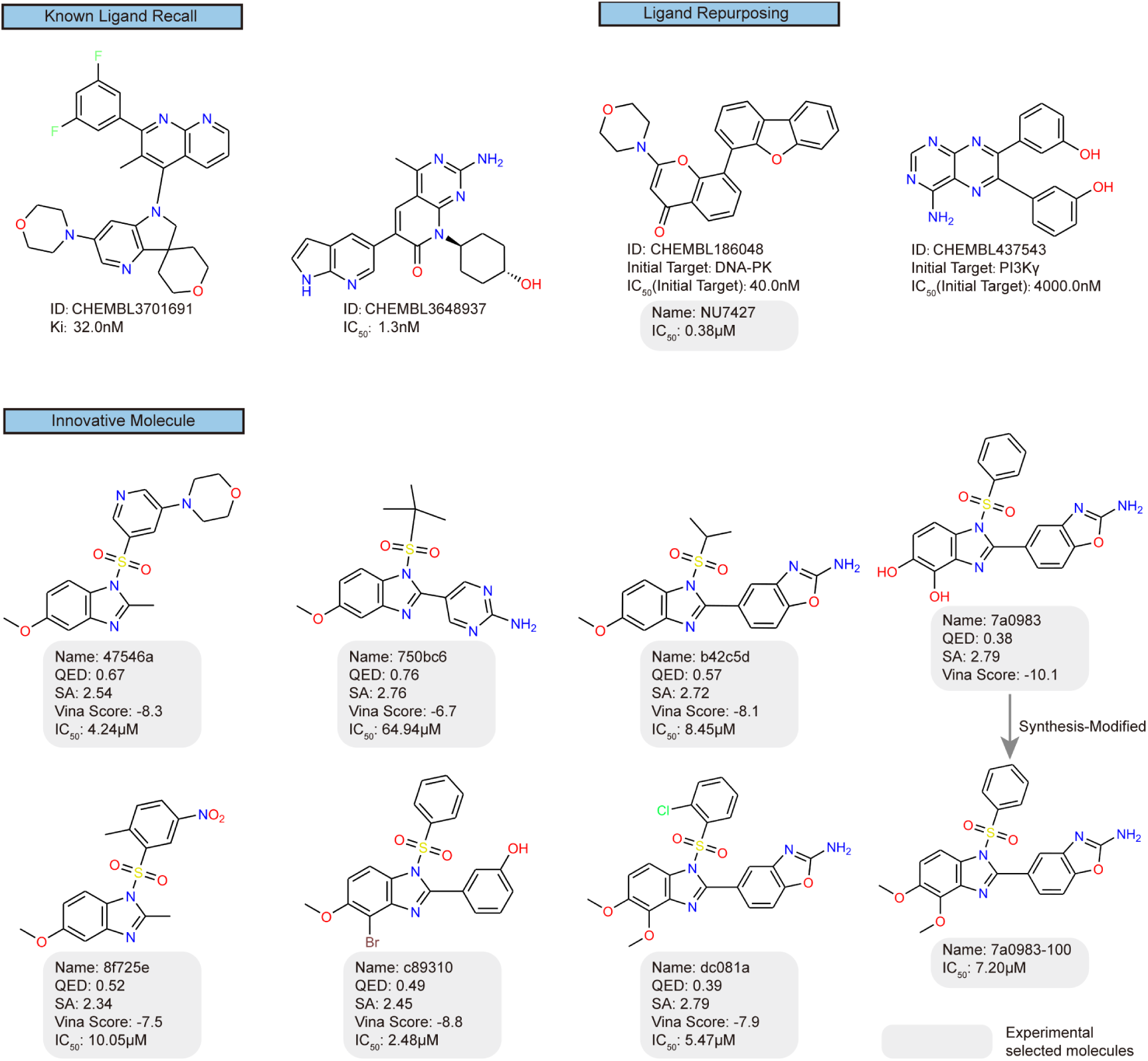
Showcase and experimental validation of molecules generated using the *de novo* option. Examples of molecules generated using the *de novo* generation of ProLigGPT, including known ligand recall, ligand repurposing, and innovative molecules. Molecules highlighted in gray boxes have undergone experimental validation, displaying their QED, SA values, Vina scores, and IC_50_ values. **a,** The successful recall of known ligands for the target protein confirms that the model has been thoroughly trained. **b,** compounds originally designed for other PI3K isoforms or different kinases were identified as potential inhibitors of PI3Kα. ProLigGPT generated the compound NU7427, which shows an IC_50_ of 40nM against DNA-PK according to BindingDB. Given that NU7427 was generated based on the PI3Kα sequence, we hypothesized it could also inhibit PI3Kα, and our experimental validation confirmed an IC_50_ of 0.38μM against PI3Kα. **c,** We selected seven molecules for experimental validation based on favorable molecular properties and docking results. Among these, six molecules were successfully synthesized. The synthesis of 7a0983 was modified to produce 7a0983-100.

**Extended Data Fig.4.**
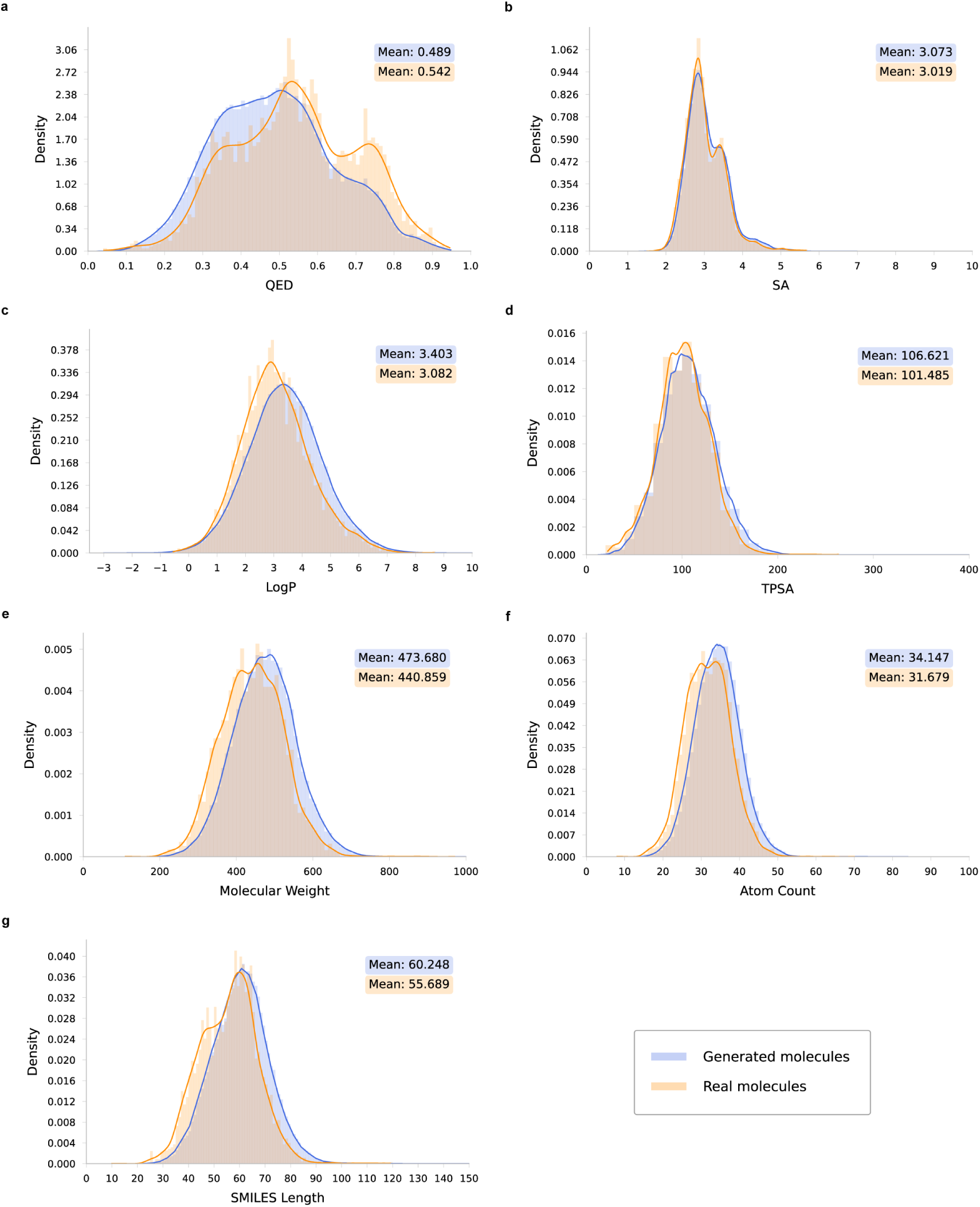
The distributions of molecular properties and metrics are similar between the generated and real molecules. **a–g**, Probability distributions of molecular properties and metrics in molecules generated by ProLigGPT and real molecules from the training dataset, including QED(**a**), SA(**b**), logP(**c**), TPSA(**d**), molecular weight(**e**), atom count(**f**), and SMILES length(**g**).

**Extended Data Fig.5.**
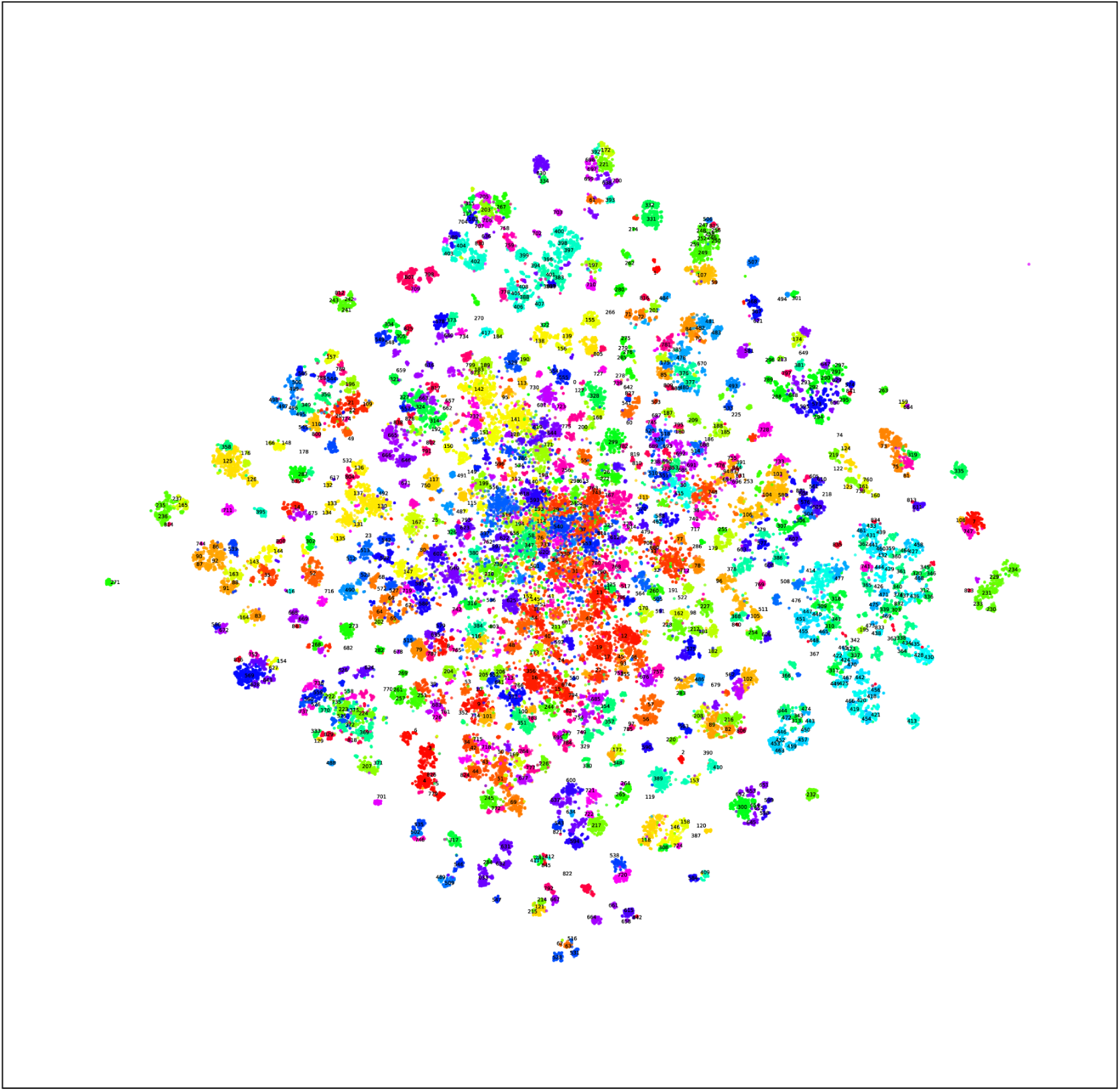
t-SNE visualization of molecular clusters by the Louvain method. The generated and reference molecules were clustered using the Louvain method and visualized via t-SNE. Each color in the diagram represents a distinct cluster, with the Louvain method identifying 846 clusters in total, thereby illustrating the relationships among the molecules. This approach effectively identifies and differentiates molecules with known and novel scaffolds, providing essential data support for subsequent molecular filtering and validation.

**Extended Data Fig.6.**
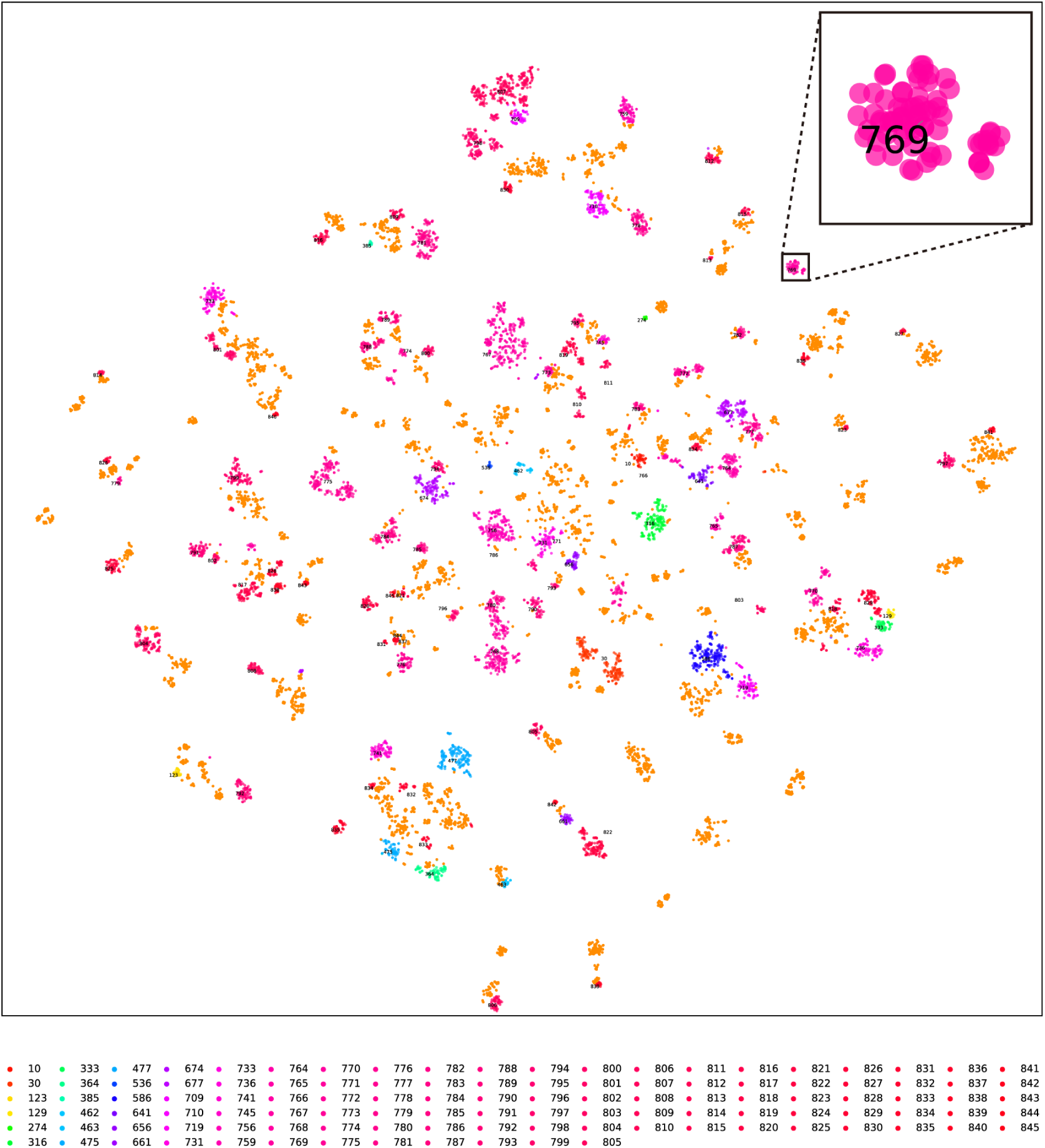
Secondary t-SNE analysis of clusters to confirm novel scaffold potential. After removing 734 clusters containing reference molecules, 112 clusters remained. The secondary t-SNE analysis of filtered clusters, excluding those with known reference molecules, indicates the potential presence of novel molecular scaffolds. Notably, Cluster 769 forms an isolated aggregation in the t-SNE plot, clearly separated from other known molecules, providing further evidence that it may contain a novel scaffold.

**Extended Data Fig.7.**
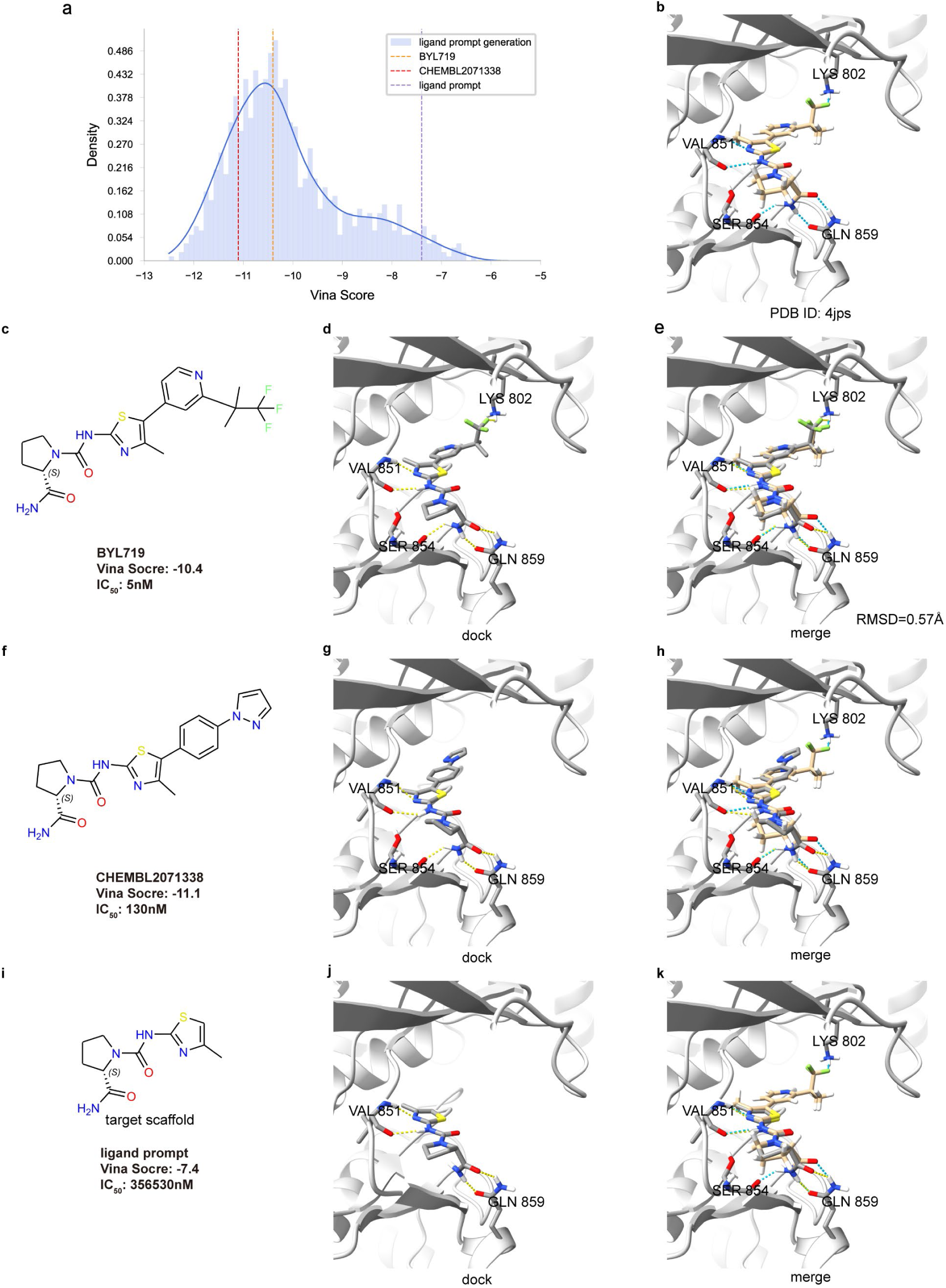
The rationale behind using the seed scaffold as a ligand prompt. **a,** Distribution of Vina score for 1000 derivative molecules generated by the ligand prompt generation. **b,** The co-crystal structure of PI3Kα with BYL719 (PDB ID: 4jps). **c,** Information about BYL719. **d,** The molecular docking result of BYL719. **e,** Merging the docking result of BYL719 with the co-crystal structure (4jps), with 0.57Å r.m.s.d.(root mean squared deviation). **f,** Information about CHEMBL2071338. **g,** The molecular docking result of CHEMBL2071338 **h,** Merging the docking result of CHEMBL2071338 with the co-crystal structure (4jps) **i,** Information about seed scaffold. **j,** The molecular docking result of the seed scaffold. **k,** Merging the docking result of the seed scaffold with the co-crystal structure (4jps).

**Extended Data Fig.8.**
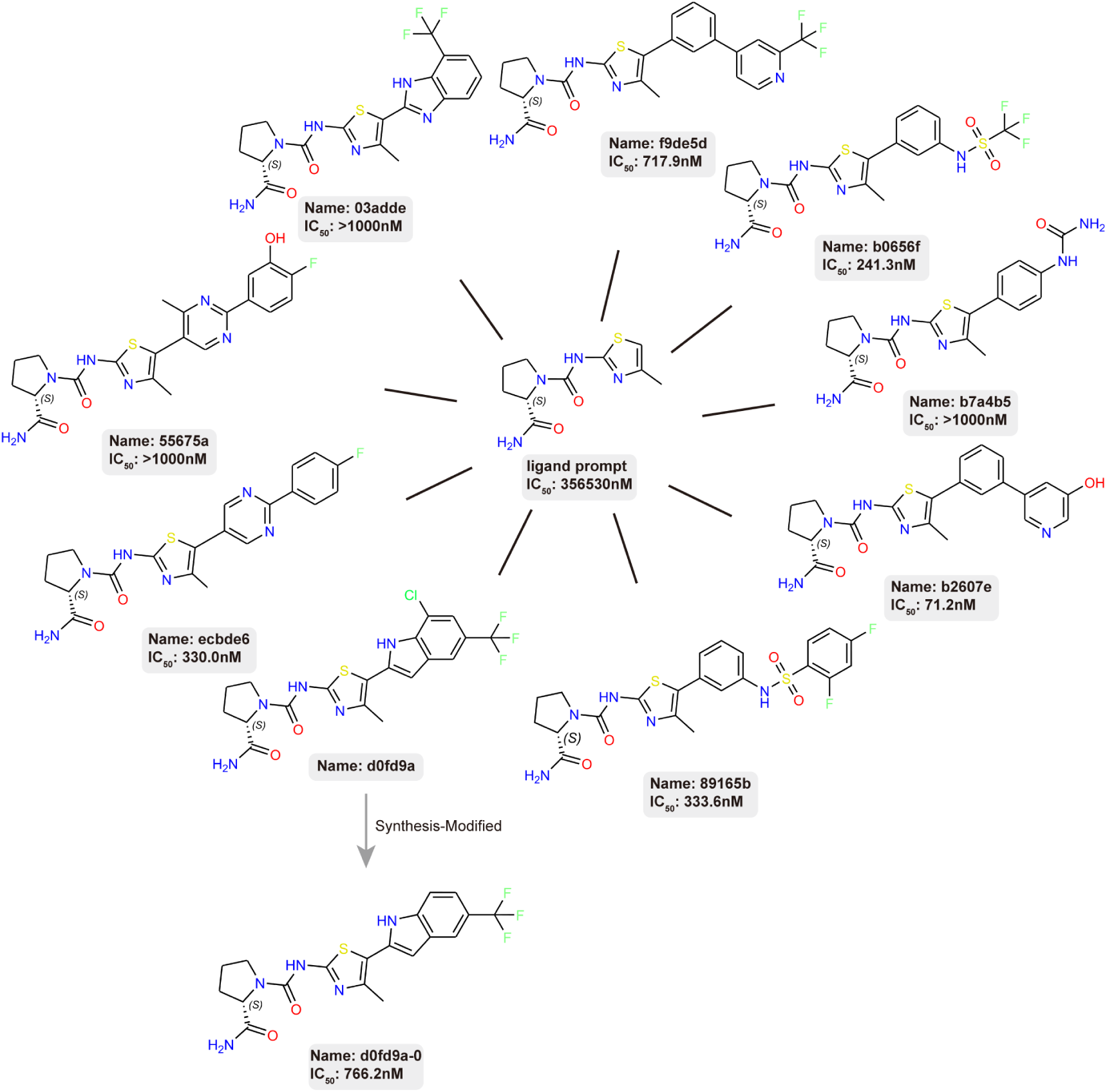
Showcase and experimental validation of molecules generated using ligand prompt option. Ligand prompt strategy enables the design of PI3Kα ligands by retaining the seed scaffold of known compounds as ligand prompts and decorating them. All the molecules shown in the figure were synthesized and evaluated through dose-response assays, with the synthesis of d0fd9a being modified to produce d0fd9a-0 due to synthetic challenges.

## Notes

### Summary of Updates

This revision updates the manuscript title from DrugGPT to ProLigGPT to better reflect its focus on protein sequence guided ligand design without structural constraints. The revised manuscript provides a more comprehensive description of the model framework, computational benchmarking, and experimental validation. It describes both de novo ligand generation and scaffold based molecular design, including experimental validation of PI3Kalpha inhibitors and a case study involving Target T1, a protein without an experimentally determined structure. To protect intellectual property, the public version withholds the identity of Target T1, selected compound structures, experimental procedures, and related figure content. The revision maintains the central research objective of developing a language model for protein targeted ligand design directly from protein sequences.

